# Gene-specific response to MuSK agonist antibody in the treatment of Congenital Myasthenic Syndromes

**DOI:** 10.1101/2025.08.29.673058

**Authors:** Kelly Ho, Ofosu Adjei-Afriyie, Ricardo Carmona-Martinez, Rohan Ray, Daniel O’Neil, Joshua Zeldin, Julien Oury, Lieselot De Clercq, Steven J. Burden, Bernhardt Vankerckhoven, Roeland Vanhauwaert, Sally Spendiff, Hanns Lochmüller

## Abstract

Congenital myasthenic syndromes (CMS) are a group of rare disorders characterized by fatigable muscle weakness and caused by impaired neuromuscular junction (NMJ) function. CMS symptoms are highly variable, but can be detrimental and lead to death. There are over 40 different genetic subtypes, including *Agrn-*CMS and *ColQ-*CMS. *Agrn* encodes for neural AGRIN, which is released from the nerve terminal and triggers muscle-specific kinase phosphorylation (pMuSK). pMuSK is essential for NMJ development and maintenance, thus AGRIN deficiency causes NMJ impairment. *ColQ* encodes for collagenous subunit Q (ColQ), which anchors acetylcholinesterase and stabilizes MuSK. As a result, ColQ deficiency results in NMJ degeneration from prolonged transmission signals and decreased pMuSK. Current treatments for *Agrn-*CMS and *ColQ-*CMS are limited, highlighting the importance of finding more efficient therapies. Recently, a MuSK agonist antibody with high affinity for the Frizzled-like domain showed remarkable rescue of a *Dok7*-CMS mouse model. We hypothesized a similar antibody could benefit *Agrn-* and *ColQ-*CMS mouse models. *Agrn-*CMS mice were treated at postnatal day 5 (P5), P15 and P35, and *ColQ-*CMS mice were treated weekly from P22 to P57. In *Agrn-*CMS mice, 3B2 treatment rescued survival, bodyweight, fibre type switching and pMuSK levels, and improved grip strength and NMJ morphology. In *ColQ-*CMS mice, 3B2 treatment was unable to rescue deficits observed. Our findings suggest that MuSK agonists may benefit patients with *Agrn*-CMS, which should be tested in clinical trials. Our study emphasizes that effective CMS treatment is gene-dependent and relies on an accurate genetic diagnosis.

## Introduction

In mice and humans, synaptic transmission at the neuromuscular junction (NMJ) involves the release of acetylcholine (ACh) from synaptic vesicles into the synaptic cleft. ACh crosses the synaptic cleft and binds to ACh receptors (AChRs) in the muscle fibre membrane. The resulting opening of AChR channels depolarizes the muscle, initiating an action potential, which leads to muscle fibre contraction. This process is rapidly terminated by acetylcholinesterase (AChE), which hydrolyzes ACh and is anchored to the basal lamina by a collagen tail (ColQ) [38].

Efficient synaptic transmission requires clustering of AChRs at the crests and upper portions of postjunctional folds, which are directly opposite to transmitter release sites, active zones, in motor nerve terminals [38]. The formation and maintenance of these pre- and postsynaptic specializations relies on the AGRIN/LRP4/MuSK pathway, reviewed in Ohno *et al* [28]. Neural AGRIN is released from motor nerve terminals and binds to low density lipoprotein receptors (LRP4) in the postsynaptic membrane. AGRIN binding promotes association between LRP4 and muscle-specific kinase (MuSK) and stimulates MuSK tyrosine phosphorylation, which is essential for AChR clustering [32, 53]. Phosphorylated MuSK (pMuSK) recruits downstream tyrosine kinase 7 (Dok7), which further aids in the phosphorylation of MuSK [2, 15]. Activated MuSK leads to tyrosine phosphorylation of AChRs, which augments the association of AChRs to receptor-associated protein of the synapse (RAPSYN) and assists in anchoring AChRs to the cytoskeleton [4, 5, 14].

Congenital myasthenic syndromes (CMS) are a diverse set of genetic neuromuscular conditions caused by impaired synaptic transmission at the NMJ. Clinically, CMS commonly presents within the first two years of life and is characterized by abnormal fatigable muscle weakness. The ocular and limb muscles are particularly affected, and the disease can be severe and even fatal with respiratory failure. Currently, mutations in over 35 genes have been shown to cause CMS [28]. *Dok7* is one of the most frequently mutated genes in CMS, and patients with *Dok7*-CMS can suffer debilitating symptoms [1, 25]. The most common disease-causing *Dok7* mutation causes truncation of the protein, leading to reduced MuSK tyrosine phosphorylation [31]. MuSK agonist antibodies (X17 and ARGX-119) provide remarkable benefit in a *Dok7-*CMS mouse model [31, 46], raising the possibility that treatment with a MuSK agonist antibody might provide similar benefit for other forms of CMS caused, at least in part, by reduced MuSK phosphorylation.

To investigate this hypothesis, we studied a CMS mouse model of AGRIN. The *Agrn^nmf380^* mouse has a point mutation that results in partial loss of function [3], where mice homozygous for this mutation are runted, display poor motor control, and usually die within a few weeks after birth [3, 40]. We also studied a second CMS mouse model lacking ColQ [7, 12, 36, 37]. Triple helical ColQ binds to AChE to assist in anchoring AChE to the synaptic basal lamina. Although loss of function mutations in *AChE* are incompatible with life, many CMS cases are caused by hypomorphic mutations in *ColQ* [11, 27, 28]*. ColQ* knockout *(ColQ*^-/-^) mice, which lack basal lamina-attached AChE, are runted, exhibit muscle weakness, and display structural abnormalities of the NMJ. *ColQ^-/-^* mice were initially reported to rarely (10-20%) survive into adulthood [12], but with extra bedding and a nutrient-soaked diet on the cage floor, *ColQ^-/-^* mice survive longer [23]. Although ColQ is not a component of the AGRIN/LRP4/MuSK pathway, it is reported to bind the Ig-like 1 and Fz domains of MuSK [29] and to stabilize MuSK at the muscle membrane [7, 29, 36, 37]. As such, we considered the possibility that boosting MuSK phosphorylation might ameliorate disease symptoms in *ColQ^-/-^* mice.

MuSK agonist antibody (ARGX-119) treatment is now undergoing clinical trials for *Dok7*-CMS (NCT06436742), so we wondered whether other forms of CMS may benefit from this treatment strategy. Our results demonstrated that 3B2, a derivative antibody of ARGX-119, fully rescued many of the cellular and phenotypic characteristics of the *Agrn^nmf380^*mouse but was ineffective in rescuing *ColQ^-/-^* mice. These findings suggest a possible therapy for *Agrn*-CMS and highlight the importance of a correct genetic diagnosis for considering treatments for CMS patients.

## Materials and methods

### Sex as a biological variable

CMS is inherited in a Mendelian fashion and thus, there are no known sex differences in prevalence, age of onset, disease severity or treatment response. A retrospective study of 235 adult CMS patients reported that 16.2% of female patients had worsened symptoms during menstruation and 32.4% of female pregnant patients had worsened symptoms during pregnancy [42]. This suggests that hormonal changes can impact CMS symptoms. Therefore, for survival, bodyweight, and motor performance, we ensured equal sexes in each group and included the first mice successfully tested from each sex.

### Animal husbandry

*Agrn^nmf380^* mice were obtained from the Burgess lab[3] and the *ColQ*^-/-^ mice [12] were given through kind donation by the Krejci laboratory (Université Paris Descartes). These mice were rederived at the University of Ottawa by the animal care and veterinary service department (ACVS) transgenic core under breeding protocol 3089. Animals were housed under 12-hour light/dark cycles and had *ad libitum* access to standard chow, water and soaked diet on the cage floor. Experimental mice were housed at the same cage locations to minimise potential confounders. Animals were weighed at least 3 times per week, and any animal found to have lost >17% of bodyweight or showing a severe phenotype as specified on our ‘in house’ monitoring sheet was humanely culled. Experimental protocol was prepared before the study commenced and is contained within a signed contract between the Children’s Hospital of Eastern Ontario Research Institute and argenx. Animal sample sizes were determined based on previous studies [40, 23] and litters were assigned at random to control or experimental conditions.

### Tissue collection

Muscle for labelling was mounted in Optimum Cutting Temperature compound (OCT) and frozen in pre-cooled isopentane before being stored at -80°C. Muscle was cryosectioned on a Leica CM1860 at 10µm and mounted on slides. Before labelling, slides were thawed at room temperature (RT) for 20 minutes. Muscle for protein analysis was snap frozen in liquid nitrogen and stored at -80°C.

### Muscle strength assessments

Mice underwent strength assessments according to standard operating protocols.[50] In the neonatal tube test, each mouse was suspended over the edge of a 50 mL Falcon tube containing bedding. The mouse was assessed for the time to fall, relative positioning of the hind limbs (hindlimb suspension score), and number of attempts the mouse made at pulling itself out of the tube. Each mouse underwent two trials with a five-minute rest period between attempts. To assess forelimb grip strength, the mouse was held by the tail and allowed to grab the grid with its forepaws before it was pulled away horizontally. After at least 10 minutes of rest, the mice were assessed for hindlimb grip strength, whereby they were held by the scruff of the neck and tail and presented to the grid at a 45° angle to allow only the hindlimbs to grip before being pulled horizontally. For both forelimb and hindlimb grip strength, each mouse was tested three times with 30 seconds of rest and the results averaged. For the hanging wire test, the mouse was acclimatized to the wire for one minute before the wire was inverted to suspend the mouse over a padded container. Time to fall was recorded and three trials were conducted with a 10-minute rest period between each trial. Mice were acclimatized to the behavioural room for at least 30 minutes before grip strength and hanging wire tests were performed.

### Muscle histology

#### Haematoxylin and eosin (H & E)

Sections were placed in tap water for 30 seconds, then into haematoxylin for 2 minutes. They were washed in tap water and placed in eosin for 30 seconds. They were washed in tap water before dehydrating through a graded ethanol series that finished with two clears using xylene. Sections were mounted with a cover slip and DPX Mountant for histology (Millipore Sigma). Tiled images were taken on a Zeiss Axio Imager at 20x magnification. Fibre area was determined using an automated system [54].

#### Myosin heavy chain (MHC) type and muscle fibre size

Sections for labelling of MHCs (fibre type) and laminin were washed in phosphate buffered saline (PBS), then blocked in 10% normal goat serum (NGS) for one hour at room temperature (RT). Primary antibodies were obtained from Developmental Studies Hybridoma Bank; BA-F8 (mouse anti-MHC1 IgG2b, 1:25), Sc-71 (mouse anti-MHC2a IgG1, 1:200), and BF-F3 (mouse anti-MHC2b, 1:200). These were applied to one section and on a second serial section, 6H1 (mouse anti-MHC2x IgGM, 1:20) was labelled. Both sections were also stained with rabbit anti-laminin IgG (1:750, Sigma, L9393). Muscles were incubated for one hour at RT and then washed in PBS. Secondary antibodies (ThermoFisherScientific): Alexa Fluor 350 IgG2b (y2b) goat anti-mouse (A-21140, 1:500), Alexa Fluor 594 IgG1 (y1) goat anti-mouse (A-21125, 1:100), Alexa Fluor 488 IgM goat anti-mouse (A-21042, 1:500), and Alexa Fluor 488 IgG goat anti-rabbit (A-11008, 1:500) were applied for one hour at RT. Following this, sections were given final washes and mounted using Vectashield hardset mounting medium (Vector Laboratories). Tiled images of the whole muscle were captured on a Zeiss Axio Imager M2 microscope. Analysis was performed with the Zeiss Zen software. In the software, fibres (>20µm) were identified using laminin and fibre MHC was determined using the histogram function for each channel.

#### NMJ labelling and analysis

Soleus muscles were washed in ice-cold PBS for 2x10 minutes and separated out into small bundles using tweezers under a stereomicroscope. They were fixed overnight at 4°C in 2% paraformaldehyde (PFA). The following morning, they were washed 2x1 hour with cold PBS. They were treated for 10 minutes with Analar Ethanol followed by 10 minutes with Analar Methanol, both at -20°C. Tissues were incubated with blocking/permeabilization solution (5% horse serum (HS), 5% bovine serum albumin (BSA), 2% Triton X-100 in PBS) for 4 hours at RT with gentle agitation. Samples were incubated with antibodies, diluted in blocking buffer (5% HS, 5% BSA), against neurofilament (mouse monoclonal IgG1, Cell Signalling, 1:100) and synaptophysin (rabbit polyclonal, ThermoFisherScientific, 1:100) overnight at 4°C with agitation and for a further 2 hours at RT the next morning. Muscles were washed in blocking buffer 4x1 hour at RT. They were incubated with Alexa 488-Conjugated α-Bungarotoxin (α-BTX) (ThermoFisherScientific, 1:250), Alexa Fluor 594 goat anti-rabbit IgG, and Alexa Fluor 594 goat anti-mouse IgG1 (ThermoFisherScientific, 1:200) for 4 hours at RT or overnight at 4°C with agitation. Samples were washed 4x1 hour in PBS and mounted using Vectashield hardset mounting medium. Images were captured using Olympus FV1000c scanning confocal microscope with FV1000 application software (FV10-ASW) software. Z-stack images at 1 μm intervals were acquired with a 63x oil immersion objective. Analysis was performed blinded according to the NMJ_Morph protocol [18].

#### Immunostaining of MuSK & pMuSK cryosections

The tissue was outlined with a PAP Pen (Sigma Z672548), dried for 30 minutes and rehydrated with PBS for 10 minutes. The tissue was fixed in 2% PFA containing a Phosphatase Inhibitor Cocktail tablet (Roche 4906845001) for 10 minutes at RT. PBS washes were performed, then the tissue was permeabilized with 0.1% Triton-X in PBS for 10 minutes at RT, followed by washing. The tissue was then incubated in blocking solution (10% goat serum + 1% BSA in PBS) for one hour. Primary antibodies, MuSK (Life Technologies PA1-1741, 1:50) and pMuSK Y755 (Abcam ab192583, 1:100) were applied overnight at 4°C in a humidified chamber. After washing, the sections were incubated with α-BTX (B13422, 1:100) and Alexa Fluor 594 Goat anti-Rabbit (A-11012, 1:100) for two hours at RT. Following PBS washes, slides were mounted with Vectashield Mounting Medium (H-1000-10), allowed to dry, and sealed with clear nail polish. Images were captured using a 40x objective on the Zeiss Axio Imager M2. Quantification of MuSK and pMuSK was conducted with Zeiss imaging software Zen Blue 3.2. A custom script was generated to semi-automatically trace α-BTX staining and provide measurements, including α-BTX area (µm²) and mean intensity values for channel AF594 (MuSK and pMuSK). Background noise from AF594 channel value was calculated and subtracted from mean intensity values. Integrated density was calculated by multiplying α-BTX area (µm²) with mean intensity values of channel AF594. An outlier test for all MuSK or pMuSK integrated density values per mouse was performed and the averages of these values were plotted.

### Quantification of pMuSK in muscle lysate through Western blot

Tibialis anterior (TA) muscles in ice-cold lysis buffer (RIPA Buffer, protease inhibitor, PhosSTOP) were homogenized using a Qiagen Tissue Lyser at 30 Hz for 3 cycles of 90 seconds, with 2 minutes of cooling on ice between cycles. The samples were rotated end-over-end at 4°C overnight. The samples were sonicated using a Sonics Vibracell Ultrasonic Processor at 20% power for 3 cycles of 3 seconds, with 2 minutes of cooling on ice between cycles. The samples were centrifuged at 16,000 g for 30 minutes at 4°C and the supernatant was collected. The protein concentration was measured using a DC protein assay (BioRad 5000112). The samples were denatured by adding Laemmli buffer (BioRad 1610747) and heated.

40 µg of protein was loaded on an 8% 1.5 M Tris-HCl gel and electrophoresed. The proteins were transferred to a pre-activated polyvinylidene difluoride (PVDF) membrane (BioRad 1620177). The membrane was blocked with EveryBlot blocking buffer (BioRad 12010020) for 15 minutes at RT. The membrane was incubated with a 1:400 dilution of anti-MuSK (phospho Y755) antibody (abcam ab192583) in EveryBlot blocking buffer for 40 hours at 4°C. The membrane was washed with tris-buffered saline 0.1% tween (TBST). The membrane was incubated with a 1:3000 dilution of goat anti-rabbit IgG secondary antibody, HRP (ThermoFisherScientific 31460) in EveryBlot blocking buffer for 1 hour at RT. After washing, the membrane was treated with Clarity Max Western ECL substrate (BioRad 1705062) for five minutes and imaged using a BioRad ChemiDoc imaging system. The membrane was washed and incubated with a 1:1000 dilution of vinculin antibody 7F9 (Santa Cruz sc-73614) in 5% milk/TBST for 1 hour at RT. After washing, the membrane was incubated with a 1:3000 dilution of goat anti-mouse IgG secondary antibody, HRP (ThermoFisherScientific 31430) in 5% milk/TBST for 1 hour at RT. After washing, bands were detected using Clarity Western ECL substrate (BioRad 1705061) and a BioRad ChemiDoc imaging system. Analysis was performed using ImageLab software.

### Quantification of MuSK in muscle lysate through ELISA

High binding 96-well half area plate (Greiner 675061) was coated with 1ug/mL anti-MuSK Ig1 clone 3B5-hIgG4-S228P (argenx) overnight at 4°C. The plate was washed with wash buffer (0.05% Tween 20 in PBS). The plate was blocked in 1% casein in 1xPBS (Bio-Rad 1610783) for 1 hour at RT, then washed. Samples were loaded onto the plate (7.5ug of protein per muscle lysate, 12-point standard curve of 10ng/mL MuSK (Biotechne 9810-MK at two-fold serial dilution) and incubated for 1 hour at RT. The plate was washed and incubated with 1ug/mL of biotinylated anti-MuSK IgG1 clone 3F6c-hIgG1-LALAdelK (argenx) for 1 hour at 22°C. The plate was washed and incubated with 20ng/mL streptavidin poly-HRP (Pierce 21140) for 30 minutes at 22°C. Following washing, total MuSK was detected with TMB substrate (Life Technologies N301). The reaction was stopped after 8 minutes with 0.5M sulfuric acid and measured at OD450-620nm with a plate reader.

### Quantification of 3B2 serum samples through ELISA

Quantification of 3B2 in serum samples was performed as described previously [46].

### Statistics

All data was collated in Excel and analysis was performed using Graphpad prism V.9.4.1. Data was first tested for normal distribution (Shapir-Wilk test for normality) and then the appropriate test performed. Normally distributed data is presented as mean ± sd, while data that does not follow a normal distribution is presented as median and interquartile range.

## Results

### Survival and bodyweight

Animals were assigned to either the drug treatment group (3B2, 3B2-hIgG1LALAdelk, the MuSK agonist antibody that binds the Fz-like domain of MuSK) or a control antibody (Mota-hIgG1LALAdelk, Mot) group, with wild type (WT) litter mates used for comparison (Supplementary Table 1). Mice assigned to 3B2 or Mot received an intraperitoneal (IP) injection on assigned days. The first IP injection was at a dosage of 20mg/kg and all subsequent injections were 10mg/kg. Animals underwent a series of behavioural assessments (neonatal tube test, grip strength, hanging wire) and blood draws. At the end of the study, mice were euthanized (CO2 followed by cervical dislocation), and tissues were weighed and collected. Due to the differing severities and presentations of the two animal models, slightly different protocols were used for each (Fig. 1A).

**Figure 1.**
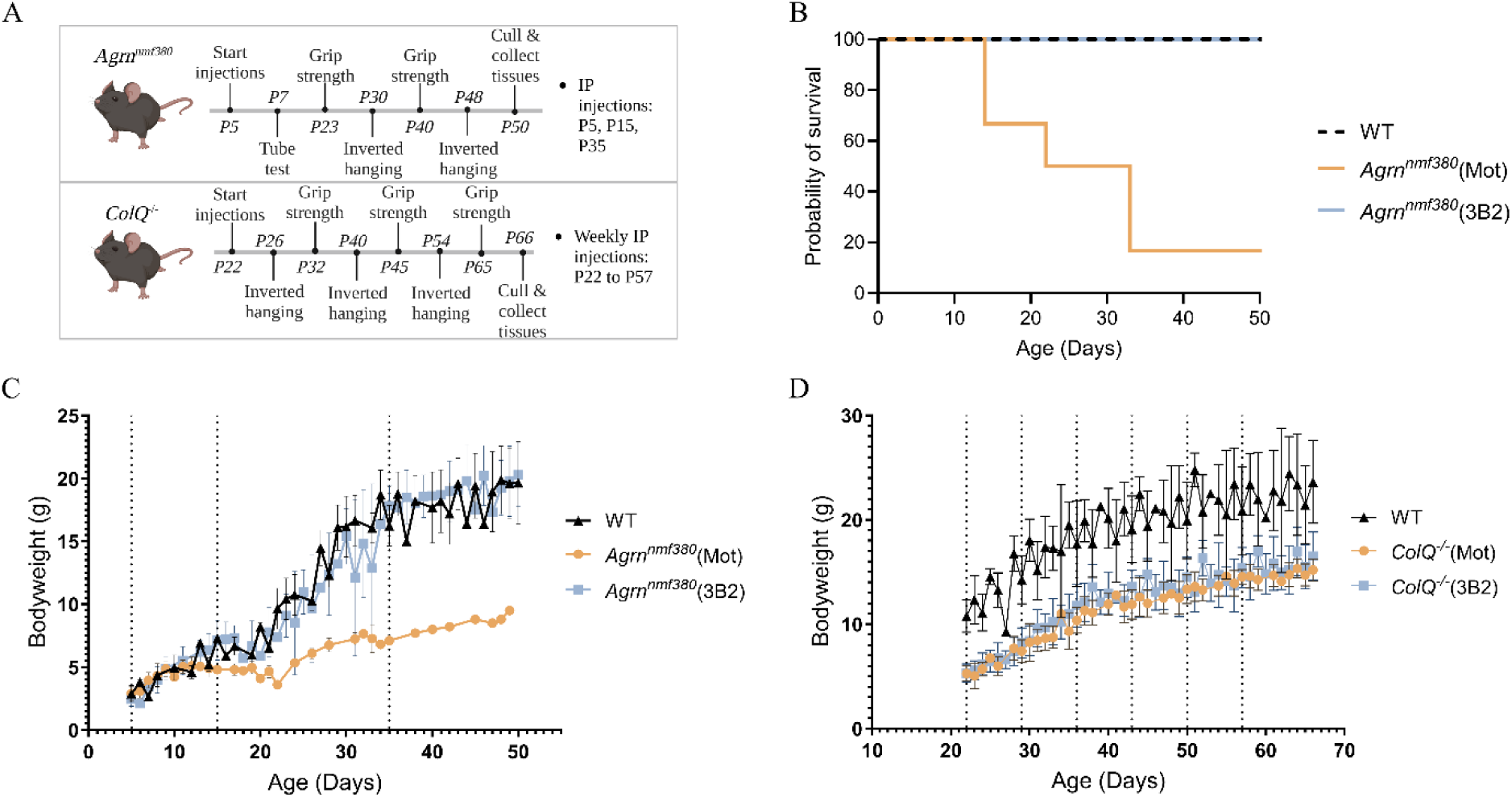
Survival and bodyweight of *Agrn^nmf380^* and *ColQ^-/-^* mice. **(A)** Treatment and testing protocol for *Agrn^nmf380^* and *ColQ*^-/-^ mice. Created with BioRender.com. **(B)** Kapplan-Meier survival showing **s**urvival of WT, *Agrn^nmf380^* (Mot), and *Agrn^nmf380^* (3B2) mice. **(C)** 3B2 treatment rescued bodyweights of *Agrn^nmf380^* mice. **(D)** 3B2 treatment did not rescue bodyweights of *ColQ^-/-^* mice. Broken vertical lines indicate IP injections. Graphs show mean ± sd. **(B)** and **(C)** n=6 mice per group. **(D)** WT n=10, *ColQ*^-/-^ (Mot) n=10, *ColQ*^-/-^ (3B2) n=12.

The *Agrn^nmf380^* study was ended when mice were P50 (Fig. 1A). ELISAs for 3B2 confirmed the accumulation of the drug in the blood of the treated mice (Supplementary Table 2). Only one *Agrn^nmf380^* (Mot) mouse survived to endpoint, and half of the mice did not survive beyond P20. In contrast, all *Agrn^nmf380^* mice treated with 3B2, like WT mice, survived to the end of the study (Fig. 1B). All *ColQ^-/-^* and WT animals survived until the end of the study (P66) (Supplementary Fig. 1), and the drug was detectible in the serum at the end of the study (Supplementary Table 3). The bodyweights of WT and *Agrn^nmf380^* (3B2) mice increased steadily over time. In contrast, the bodyweights of the few surviving *Agrn^nmf380^* (Mot) mice plateaued after 15 days and increased only slowly thereafter (Fig. 1C). Unlike the *Agrn^nmf380^* mice, 3B2 did not rescue the loss of bodyweight of *ColQ*^-/-^ mice (Fig. 1D).

### Muscle strength

*Agrn^nmf380^* mice were subjected to hindlimb suspension, grip strength and inverted screen tests throughout the study (Fig. 2A). We measured muscle strength of *Agrn^nmf380^* mice two days following the first injection at P7 using the hindlimb suspension test (Fig. 2B, 2C & Supplementary Fig. 2). *Agrn^nmf380^* (3B2) mice held on with their hindlegs for longer than *Agrn^nmf380^* (Mot) mice (Fig. 2B).

**Figure 2.**
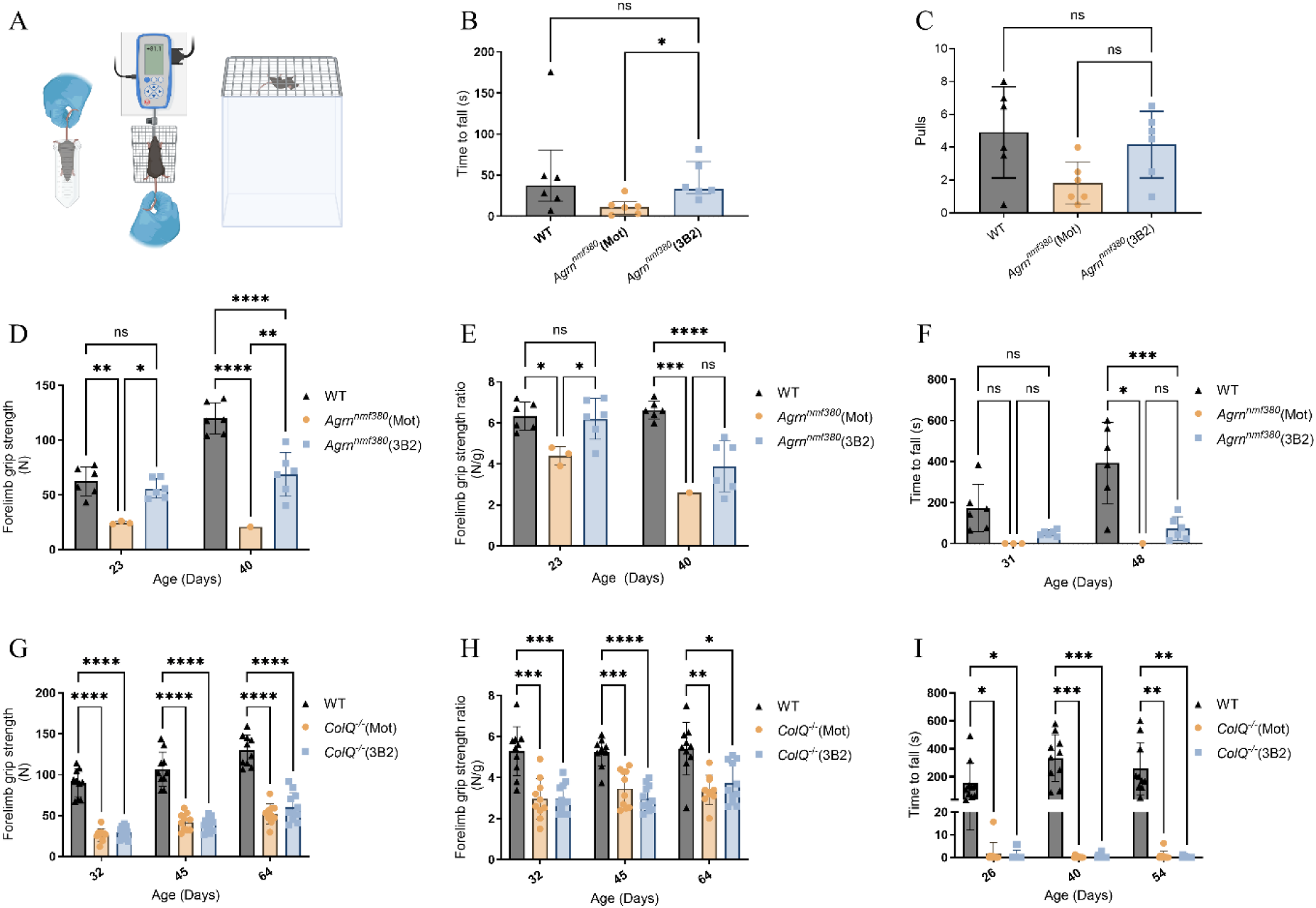
Motor behavioural testing of *Agrn^nmf380^* and *ColQ^-/-^* mice. **(A)** Schematic illustration of hindlimb suspension test, grip strength test and inverted screen test. Created with BioRender.com. **(B)** *Agrn^nmf380^* (3B2) mice held on for longer than *Agrn^nmf380^* (Mot) mice in the hindlimb suspension test at P7-9. Median and IQR, Kruskal-Wallis test with Dunn’s correction for multiple comparisons. **(C)** There were no significant differences in the number of pulls in the hindlimb suspension test. Mean ± sd. 1-way ANOVA with Tukey’s correction. **(D)** 3B2 treatment improved grip strength of *Agrn^nmf380^* animals at P23 and P40. **(E)** After normalization for bodyweight, 3B2 treatment improved grip strength of *Agrn^nmf380^*animals at P23. **(F)** Inverted screen test of WT, *Agrn^nmf380^* (Mot) and *Agrn^nmf380^*(3B2) mice. **(G)** and **(H)** 3B2 treatment did not improve grip strength of *ColQ*^-/-^ mice. **(I)** 3B2 treatment did not improve ability to hold on in inverted screen test of *ColQ*^-/-^ mice. Graphs show mean ± sd. 2-Way ANOVA with Tukey’s multiple comparisons correction. **(A)** and **(B)** n=6 animals per group **(C)** to **(E)** to At P23 & 31: WT n=6, *Agrn^nmf380^*(Mot) n=3, *Agrn^nmf380^* (3B2) n=6. At P42 & P48: WT n=6, *Agrn^nmf380^*(Mot) n=1, *Agrn^nmf380^* (3B2) n=6 **(F)** to **(H)** WT n=10, *ColQ*^-/-^ (Mot) n=10, *ColQ*^-/-^ (3B2) n=12. *p<0.05, **p<0.005, ***p<0.001, ****p<0.0001, ns=non-significant.

Once mice were weaned, they underwent strength testing using the grip strength meter and inverted screen test. It should be noted that these assessments for *Agrn^nmf380^* (Mot) mice were impacted by the poor survival of these mice, resulting in a low n for this condition. Nonetheless, the grip strength of surviving *Agrn^nmf380^* (Mot) mice was reduced by two- to six-fold compared to WT mice at P23 and P42, respectively (Fig. 2D & 2E). In contrast, *Agrn^nmf380^* (3B2) mice had WT-levels of grip strength at P23 (Fig. 2D). Although this rescue was not sustained, as their grip strength was less (approximately 2-fold) than WT mice at P42. A similar pattern was observed for hindlimb grip strength (Supplementary Fig. 3A & 3B).

At P31, there were no significant differences in inverted screen test performance between WT, *Agrn^nmf380^* (Mot) and *Agrn^nmf380^* (3B2) mice due to the large variability in data. By P48, WT mice held on longer than both *Agrn^nmf380^*(Mot) and *Agrn^nmf380^* (3B2) mice (Fig. 2F). Although there is a lack of significance in the holding times between *Agrn^nmf380^* (Mot) and *Agrn^nmf380^* (3B2) mice, from a functional benefit angle, the differences are drastic. At P31, *Agrn^nmf380^* (Mot) held on for an average of 0 seconds, while *Agrn^nmf380^* (3B2) mice held on for an average of 51.4 seconds. At P48, the one surviving *Agrn^nmf380^* (Mot) mouse held on for an average of 1.3 seconds, while *Agrn^nmf380^* (3B2) mice held on for an average of 72.3 seconds (Fig. 2F). While not in our original statistical plan, a post-hoc analysis classifying mice as ‘non-holders’ (<2s) and holders (>2s) showed that there was a significant improvement in treated animals at P31 (Supplementary Table 4). It was not possible to examine this at P48 due to the low n number of *Agrn^nmf380^* (Mot) mice. Again, similar patterns were observed when the data was normalised for bodyweight (Supplementary Fig. 3C). Like the *Agrn^nmf380^* mice, *ColQ^-/-^*mice were significantly weaker than their WT littermates in forelimb grip strength (Fig. 2G). This difference remained across the 44 day study and following normalisation for bodyweight (Fig. 2H). Unlike the *Agrn^nmf380^* mice, there was no rescue of forelimb grip strength in *ColQ^-/-^* mice animals with 3B2 treatment. While hindlimb strength was significantly higher in WT mice compared to their *ColQ^-/-^* counterparts, this difference was not maintained once values were normalised for bodyweight. No rescue was observed with 3B2 treatment (Supplementary Fig. 3D & 3E). On the inverted hanging wire test, WT animals held on for significantly longer than *ColQ^-/-^* mice before and after normalisation for bodyweight (Fig. 2I & Supplementary Fig. 3F). Similarly to grip strength, there was no rescue with 3B2 treatment in the *ColQ^-/-^*mice.

### Muscle characteristics

At the end of the experiments, muscle tissue was collected and weighed (Supplementary Fig. 4A & 4B). When calculated relative to bodyweight, 3B2 treatment rescued muscle weight to WT levels (Fig. 3A). Both WT and *Agrn^nmf380^* (3B2) animals had significantly heavier quadriceps (Quad) and gastrocnemius (Gas) muscles than *Agrn^nmf380^* (Mot) animals (Fig. 3A). The difference in muscle weight normalised to bodyweight suggested that the increased muscle weight cannot be fully explained by variations in age or overall bodyweight of the animals. Following normalisation for bodyweight, 3B2 treated *ColQ^-/-^* mice had a partial rescue in quad muscle weight than *ColQ^-/-^* (Mot) mice (Fig. 3B).

**Figure 3.**
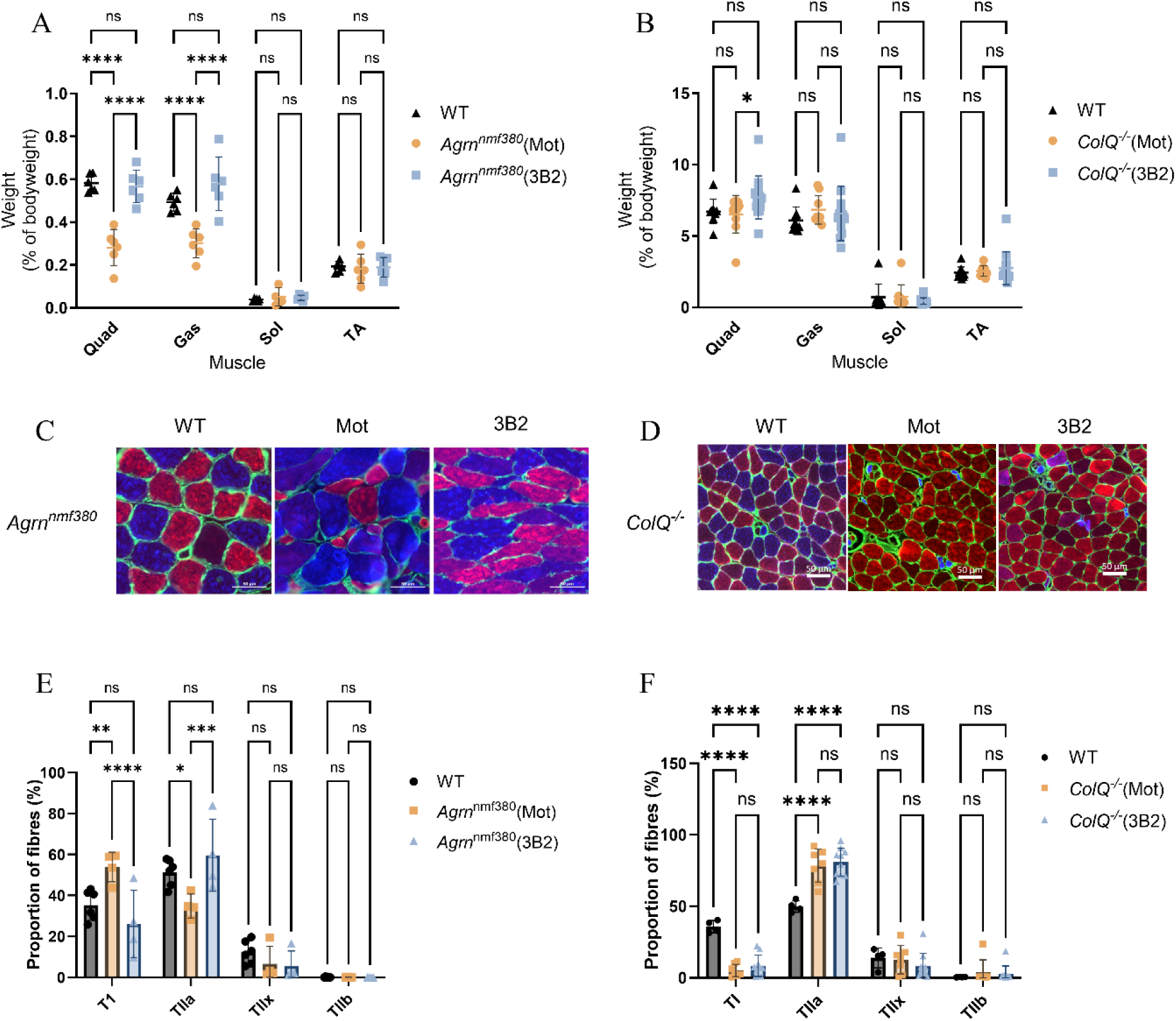
Muscle characteristics of *Agrn^nmf380^* and *ColQ^-/-^* mice. Quadriceps (Quad), gastrocnemius (Gas), soleus (Sol), and tibialis anterior (TA) muscles were weighed and normalised to bodyweight. (A) *Agrn^nmf380^* (3B2) mice had Quad and Gas muscle weights indistinguishable from WT. (B) *ColQ^-/-^* (3B2) mice had heavier Quad muscles compared to both *ColQ^-/-^* (Mot) and WT mice. (C) & (D) Cross sections of Sol muscles were stained to determine MHC type. Blue = type 1 (TI), red = type 2a (TIIa), green = type 2x (TIIx & laminin), no labelling = type 2b (TIIb). (E) *Agrn^nmf380^* (3B2) mice demonstrated a full rescue of the fibre type switching observed in *Agrn^nmf380^* (Mot) mice. (F) 3B2 treatment did not reverse the slow to fast MHC fibre type switching observed in *ColQ^-/-^* (Mot) mice. Graphs show mean ± sd. 2-Way ANOVA with Tukey’s multiple comparisons correction. (A) n=6 animals per group (B) WT n=10, *ColQ*^-/-^ (Mot) n=10, *ColQ*^-/-^ (3B2) n=12 (E) WT n=6, *Agrn^nmf380^*(Mot) n=4 (Unequal sexes), *Agrn^nmf380^* (3B2) n=4. *p<0.05, **p<0.005, ***p<0.001, ****p<0.0001.

We examined myosin heavy chain (MHC) type expression in the soleus muscles of *Agrn^nmf380^* and *ColQ^-/-^* mice (Fig. 3C & 3D). *Agrn^nmf380^*mice exhibited a fast to slow fibre type shift (Fig. 3E), as previously described in this model and in patients [40, 52]. This abnormality was reversed in *Agrn^nmf380^* mice treated with 3B2 (Fig. 3E). As previously shown [23, 36], *ColQ*^-/-^ mice had considerably fewer type 1 (TI) slow fibres when compared to WT, following a slow to fast fibre type shift, which was not reversed with 3B2 treatment (Fig. 3F).

WT mice had significantly larger TI fibres than *Agrn^nmf380^*(Mot) mice (Table 1). The reduction in muscle fibre size was rescued in *Agrn^nmf38^* mice by 3B2 treatment. The largest differences in fibre size were seen in the faster type 2 (TII) MHC types, as *Agrn^nmf380^*(Mot) mice had smaller TIIa and TIIx fibres than WT animals. Treatment with 3B2 rescued this deficit, consistent with the rescue in fibre type switching (Fig. 3E). This improvement in fibre size with 3B2 treatment was also seen in the quadriceps muscle with H & E histochemistry (Supplementary Fig. 5A & 5B). As with *Agrn^nmf380^* mice, WT mice had significantly larger muscle fibres than *ColQ^-/-^* mice; however, 3B2 failed to increase myofibres size (Table 1).

**Table 1:**
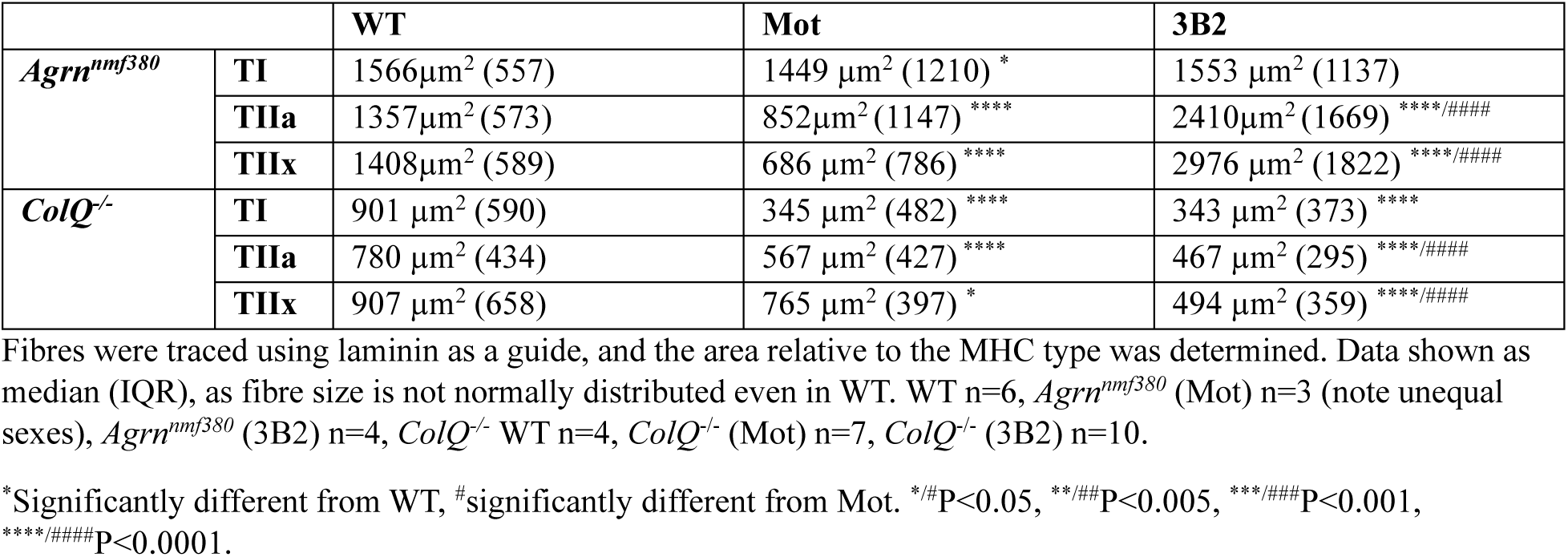
Fibre area by MHC type.

### NMJ structure

To determine changes in NMJ morphology in the soleus muscle, analysis was performed on maximum intensity projections of NMJs taken using confocal microscopy (Fig. 4A) and analysed as previously described[18, 40] (Supplementary Table 5 & 6). There were many differences between WT and *Agrn^nmf380^*(Mot) mice, especially among the presynaptic variables. Administration of 3B2 rescued most of the presynaptic variables, including nerve terminal perimeter (Fig. 4B) and complexity (Fig. 4C). While every effort was made to capture all NMJs in the muscle, there were fewer visible in the *Agrn^nmf380^* (Mot) mice. In addition, a greater number of these NMJs had minimal postsynaptic labelling visible (50% in *Agrn^nmf380^*(Mot) vs <20% in WT and *Agrn^nmf380^* (3B2) mice, (Supplementary Table 5). Nonetheless, *Agrn^nmf380^* (3B2) mice had a larger endplate area than *Agrn^nmf380^* (Mot) mice (Fig. 4D). Interestingly, *Agrn^nmf380^* (3B2) mice had the highest degree of fragmentation of all three groups, which was reflected in their lowest value for ‘average area of AChR clusters’ (Supplementary Table 5). *ColQ^-/-^* mice also had reductions in many morphological features of the NMJ (Supplementary Table 6), which were not rescued by 3B2 treatment. Additional morphological measures of the NMJ were further reduced in *ColQ*^-/-^ (3B2) mice (Fig. 4E to 4G), in contrast to *Agrn^nmf380^*mice.

**Figure 4.**
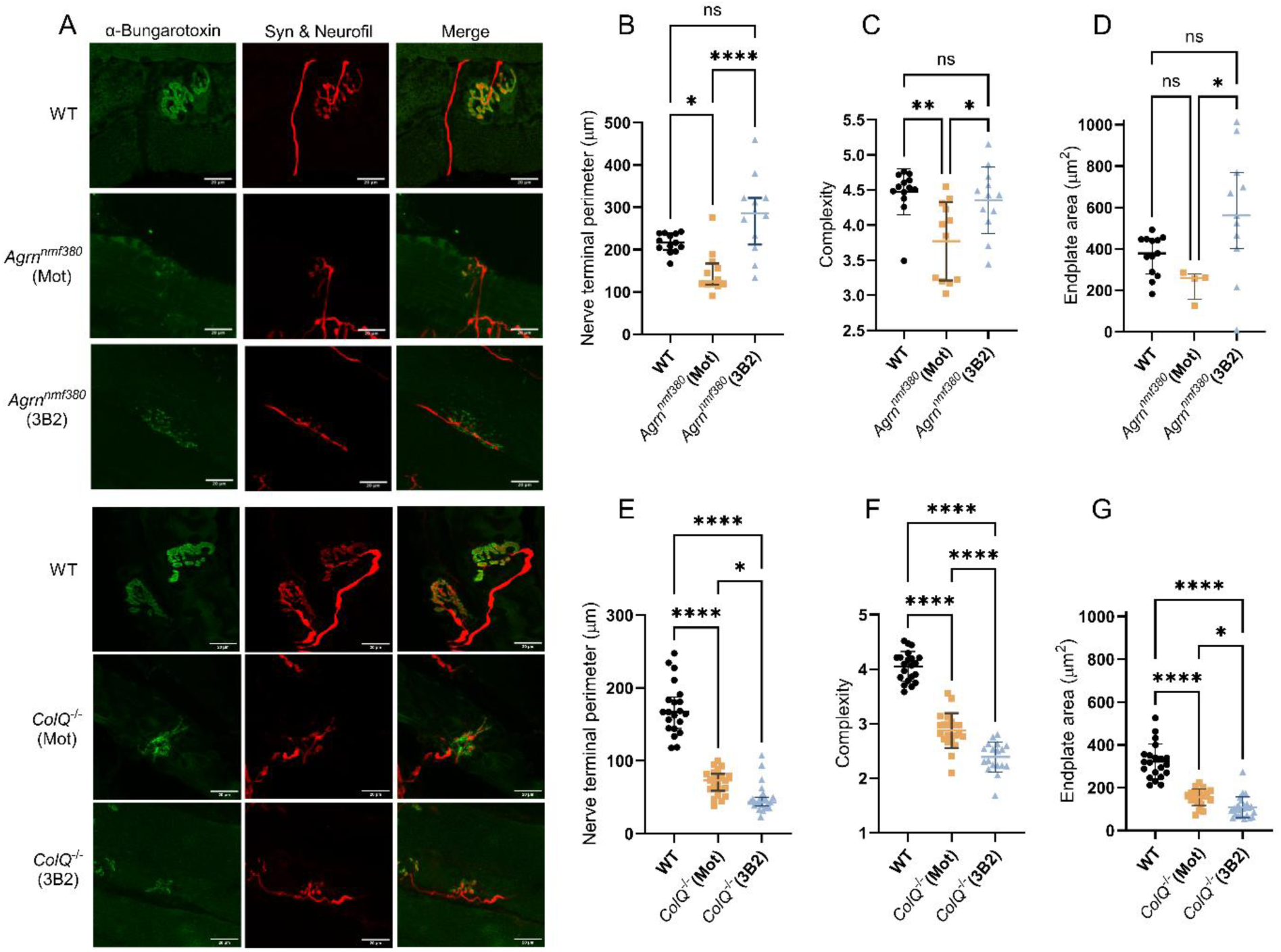
NMJ morphological analysis of *Agrn^nmf380^* and *ColQ^-/-^* mice. (A) NMJs in whole soleus muscles were labelled using α-Bungarotoxin (green), synaptophysin (Syn, red), and neurofilament (Neurofil, red). Maximum intensity projection images were analysed to calculate various morphological features of the NMJs. Scale bar is 20 µm. 3B2 treatment rescued (B) nerve terminal perimeter, (C) complexity and (D) endplate area in *Agrn^nmf380^* mice. 28-72 NMJs were analysed per group. 3B2 treatment further reduced (E) nerve terminal perimeter, (F) complexity and (G) endplate area in *ColQ^-/-^* mice. 122-127 NMJs were analysed per group. (B), (D), (E) & (G) Graphs show median ± IQR, Kruskal-Wallis test followed by Dunn’s multiple comparisons test. (C) & (F) Graphs show mean ± sd. 1-way ANOVA with Tukey’s multiple comparisons correction. (B) to (D) n=6 animals per group. (E) to (G) WT n=7, *Agrn^nmf380^*(Mot) n=3, *Agrn^nmf380^* (3B2) n=5. *p<0.05, **p<0.005, ****p<0.0001.

### Quantification of MuSK and phosphorylated MuSK

MuSK agonist antibodies that bind the Fz-like domain of MuSK stimulate MuSK dimerization and phosphorylation *in vivo* [31]. We investigated the effects of 3B2 treatment on MuSK and pMuSK in two ways. Firstly, we performed immunofluorescence labelling at the NMJ in the gastrocnemius muscle (Fig. 5A). In *Agrn^nmf380^* animals, there was no reduction in MuSK expression at the NMJ, nor a change with 3B2 treatment (Fig. 5B). There was, however, a reduction in pMuSK levels in *Agrn^nmf380^* mice that was rescued by 3B2 treatment (Fig. 5C). When the ratio of pMuSK/MuSK was determined, there was a 4.3-fold increase in *Agrn^nmf380^*(3B2) mice (Fig. 5D). We also examined pMuSK levels in TA muscle lysates using Western blot (Fig. 5E & Supplementary Fig. 6). *Agrn^nmf380^* (Mot) mice had a 0.1-fold decrease in pMuSK compared to WT, which was rescued with 3B2 treatment (Fig. 5F). We quantified total MuSK levels in the TA with an enzyme-linked immunosorbent assay (ELISA) and found an increase in MuSK in *Agrn^nmf380^* (Mot) mice (Fig. 5G), consistent with increased MuSK expression in denervated, inactive muscles [45].

**Figure 5.**
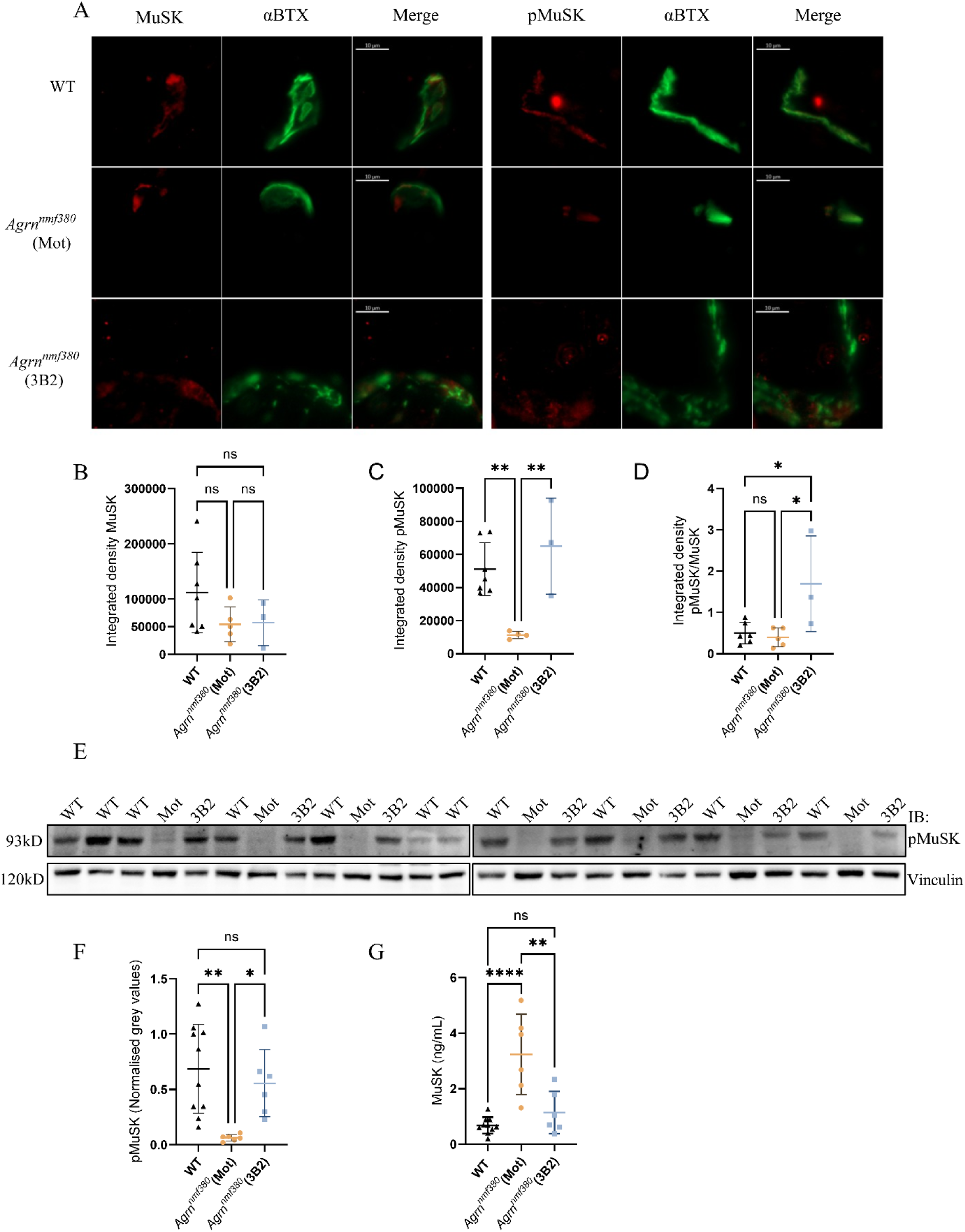
MuSK and pMuSK in *Agrn^nmf380^*mice. **(A)** Gastrocnemius muscles were labelled with α-bungarotoxin (α-BTX), MuSK, and pMuSK, and the integrated density determined. **(B)** No differences in MuSK at the NMJ were detected between groups. **(C)** 3B2 treatment increased pMuSK and **(D)** and pMuSK/MuSK at the NMJ in *Agrn^nmf380^* mice. **(E)** Western blot of TA lysates from WT, *Agrn^nmf380^* (Mot) and *Agrn^nmf380^* (3B2) mice. Note the number of bands on the image does not match the number of data points as some samples were run on both gels to allow normalisation across two blots. **(F)** pMuSK levels were reduced in *Agrn^nmf380^* (Mot) mice, which was completely reversed with 3B2 treatment. **(G)** When quantified with an ELISA, *Agrn^nmf380^* (Mot) mice had higher total MuSK levels in TA muscle lysate. Graphs show mean ± sd. Tukey’s multiple comparisons correction. **(B)** to **(D)** WT n=7, *Agrn^nmf380^* (Mot) n=5, *Agrn^nmf380^* (3B2) n=3 **(F)** WT n=10, *Agrn^nmf380^* (Mot) n=7, *Agrn^nmf380^* (3B2) n=7 *p<0.05, **p<0.005, ****p<0.0001.

In *ColQ^-/-^* mice, we likewise performed immunofluorescent labelling of MuSK and pMuSK at the NMJ in the gastrocnemius muscle (Fig. 6A). *ColQ^-/-^* mice had a 0.6-fold decrease in MuSK compared to WT mice, which was rescued to WT levels with 3B2 treatment (Fig. 6B). There was no reduction in pMuSK (Fig. 6C) or the pMuSK/MuSK ratio (Fig. 6D) in *ColQ*^-/-^ mice. Western Blot of the TA lysate (Fig. 6E & Supplementary Fig. 7) showed that *ColQ*^-/-^ (Mot) mice had less pMuSK than WT (Fig. 6F). ELISA quantification of total MuSK showed that there was an increase in total MuSK in *ColQ*^-/-^ (3B2) mice when compared to both WT and *ColQ*^-/-^ (Mot) mice (Fig. 6G).

**Figure 6.**
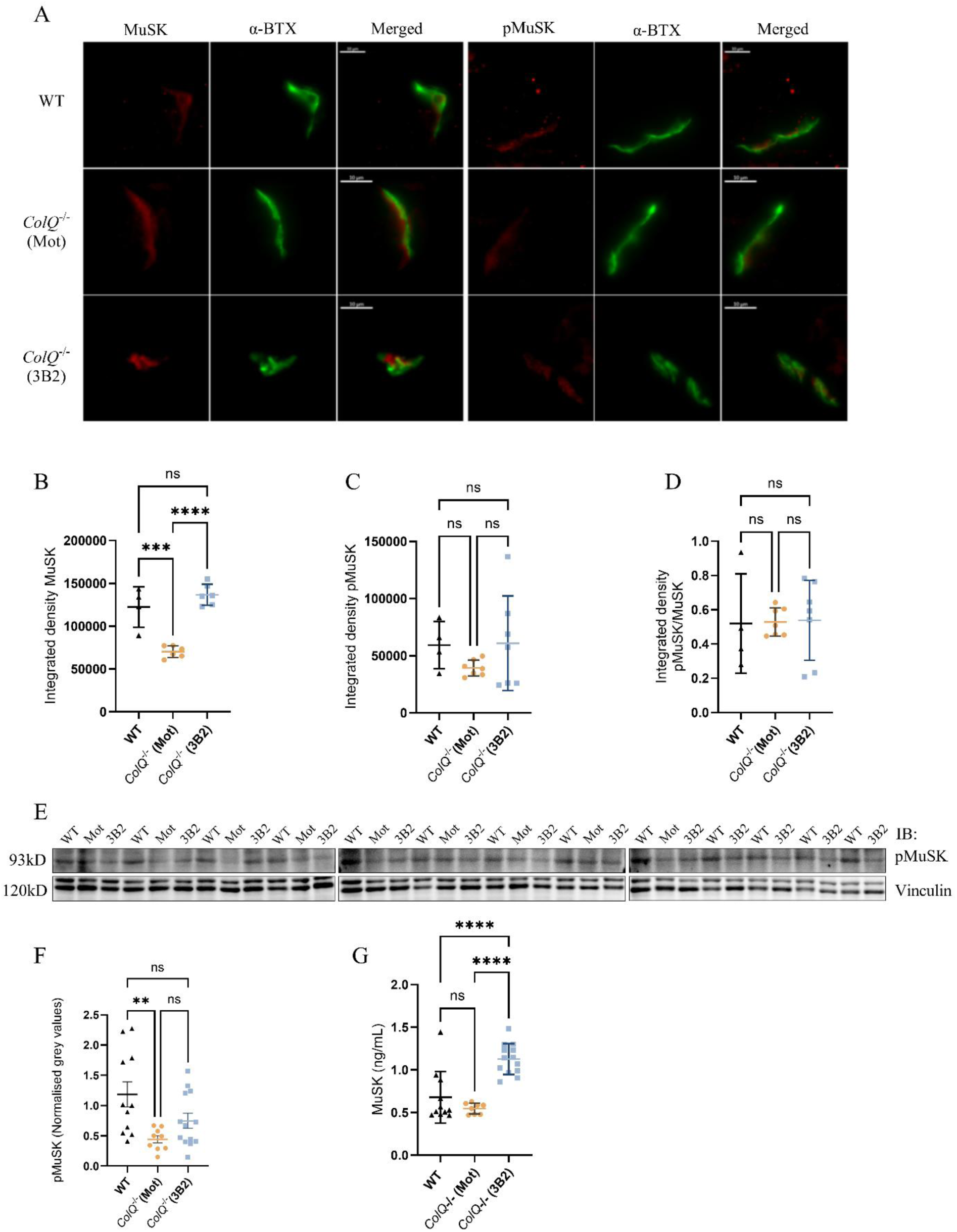
MuSK and pMuSK in *ColQ*^-/-^ mice. **(A)** Gastrocnemius muscles were labelled with α-bungarotoxin (α-BTX), MuSK, and pMuSK, and the integrated density determined. **(B)** 3B2 treatment rescued total MuSK levels at the NMJ in *ColQ*^-/-^ mice. 3B2 treatment did not increase **(C)** pMuSK or **(D)** pMuSK/MuSK ratio at the NMJ in *ColQ*^-/-^ mice. **(D)** Western blot of TA lysates from WT, *ColQ*^-/-^ (Mot) and *ColQ*^-/-^ (3B2) mice. WT n=11, *ColQ*^-/-^ (Mot) n=9, *ColQ*^-/-^ (3B2) n=13 **(F)** *ColQ*^-/-^ (Mot) mice had significantly less pMuSK than WT animals but no significant increase was observed with 3B2 addition. Note the number of bands on the image does not match the number of data points as some samples were run on both gels to allow normalisation across three blots. **(G)** MuSK was quantified in TA muscles with an ELISA. WT n=11, *ColQ*^-/-^ (Mot) n=8, *ColQ*^-/-^ (3B2) n=13 **(G**) Graphs show mean ± sd. 1-way ANOVA with Tukey’s multiple comparisons correction. **(B)** to **(D)** WT n=4, *ColQ*^-/-^ (Mot) n=6, *ColQ*^-/-^ (3B2) n=6. (F) WT n=11, *ColQ*^-/-^ (Mot) n=8, *ColQ*^-/-^ (3B2) n=13 **p<0.005, ***p<0.001, ****p<0.0001.

## Discussion

CMS are a group of genetic disorders in which mutations in genes encoding proteins at the NMJ impair neuromuscular transmission. *Agrn-*CMS and *ColQ-*CMS patients have abnormal NMJ morphology and function, presenting with fatigable muscle weakness. The most common CMS treatment, acetylcholinesterase inhibitors, are ineffective and sometimes detrimental for both *Agrn-* and *ColQ-*CMS [43]. MuSK agonist antibodies that target the Fz-like domain and stimulate MuSK phosphorylation have improved survival, weight gain, NMJ formation and maturation, and completely rescued motor function when assessed with behavioural tests in Dok7-CMS mice. In some cases, these treated mice have proceeded to become fertile adults [31, 46].

This is the first time an agonist antibody treatment has been shown to rescue the phenotype in *Agrn-*CMS. Coupled with an easy delivery system, we hypothesize that 3B2 may be a therapeutic option for patients with *Agrn*-CMS. *Agrn^nmf380^* mice have a partial loss of function of AGRIN, resulting in reduced survival, bodyweight, and motor strength when assessed through grip strength and inverted screen tests. We and others have reported that untreated *Agrn*^nmf380^ mice often die from P12 onwards [3, 40]. The remarkable rescues in survival and bodyweight in our 3B2-treated *Agrn^nmf380^* mice were consistent with the results in *Dok7-*CMS mice with Fz-like MuSK agonist antibody treatment. 3B2 treatment increased time to fall in the hindlimb suspension test and grip strength in *Agrn^nmf380^* mice. However, they did not experience a full rescue of grip strength back to WT levels at the later time point of P40. This may be indicative that more frequent dosing of 3B2 is required in the *Agrn*^nmf380^ mice to maintain WT level grip strength. In the inverted screen test, 3B2 treatment did not significantly improve latency to fall or holding impulse. However, the differences we saw in *Agrn^nmf380^* mice with 3B2 treatment (0 seconds to 51.4 seconds) were an extremely biologically relevant improvement. When a post-hoc analysis was performed by classifying mice as ‘non-holders’ (<2s) and holders (>2s), 3B2 treatment significantly increased the number of *Agrn^nmf380^* mice classified as holders at P31. Future studies may benefit by analyzing this test in groups of non-holders, holders, and extended holders, to avoid missing what may be relevant results. In addition, beginning treatment even earlier could result in more beneficial effects.

An increase in muscle weight of *Agrn^nmf380^* (3B2) mice back to WT levels is not surprising given their increase in bodyweight; this improvement remained after normalizing to bodyweight, suggesting that strength gains are also due to an improvement intrinsic to the nerve-muscle complex. *Agrn^nmf380^*mice have smaller muscle fibres [40] and smaller TI, TIIa, and TIIx fibres. This difference was more pronounced in the TIIa and TIIx fibres, which is in line with the fast to slow fibre type shift we observed in these animals. These differences were completely reversed in the *Agrn^nmf380^*(3B2) mice, and in the case of the TIIa and TIIx fibres, an increase in size compared to WT mice occurred. While we have previously observed rescues of these features in *Agrn^nmf380^*animals with treatment of a modified form of AGRIN [40], this is the first time we have seen a recovery back to WT levels. The MHC type is influenced by several factors including neural input, in the case of NMJ dysfunction this is perturbed, and fibre-type conversion or a shift to an undifferentiated state can be observed. The rescue of this with 3B2 suggests an improvement in the neural input secondary to NMJ improvement. The mosaicism of MHC types in a muscle is important to preserve energy efficiency, adaptability, precision and control of movement [33].

In previous studies [31, 46], it was shown that MuSK agonist antibodies were able to rescue synapse formation and maturation in the *Dok7-*CMS mouse. In the case of *Agrn^nmf380^* (3B2) mice, we observed extensive rescue of morphology in both pre- and postsynaptic components of the NMJ. While the ability of 3B2 to stimulate the postsynaptic apparatus is in line with its proposed mechanism of action, the indirect effects on the presynaptic side are remarkable. While it is possible that presynaptic changes are secondary to a healthier muscle, it is also possible that a more direct mechanism is contributing to these improvements. Previous research has shown the ability of the AGRIN/LRP4/MuSK pathway to influence the presynapse [16], with *Agrn* mutations negatively impacting the innervating axon [3] and AGRIN supplementation also being able to improve the presynaptic morphology [21, 40]. Other pathway components that have been shown to impact the presynaptic terminal include MuSK [19, 47] and LRP4 [8, 35, 51].

NMJ fragmentation is often used as an indicator of NMJ pathology. However, in this and a previous study [40], an improvement in phenotype and muscle characteristics was accompanied by an increase in NMJ fragmentation. Previous research has also shown that fragmentation is not necessarily associated with impaired transmission [49] and may indeed be a method of compensation to restoration of function [39].

Both Western blot and immunofluorescent labelling showed *Agrn^nmf380^* (Mot) mice had severely reduced pMuSK levels, which were rescued with 3B2 treatment. Conversely, *Agrn*^nmf380^ (Mot) mice had higher total MuSK levels than both *Agrn^nmf380^* (3B2) and WT animals. MuSK expression is dramatically induced in adult muscle following denervation [45], which has been previously reported in *Agrn^nmf380^*mice [3] and in *Agrn-*CMS patients [17].

Alternatively, these high levels could be due to the young age at which *Agrn*^nmf380^ (Mot) mice died. During the first weeks of postnatal development, the NMJ undergoes intensive maturation, and MuSK is highly involved in the prepatterning of muscles, while in adult mice, expression is limited to the subsynaptic nuclei in the central region of the muscle [5, 19]. MuSK expression has been shown to dramatically downregulate shortly after birth in rats [45]. While ColQ does not sit in the AGRIN/LRP4/MuSK pathway, it has been reported to bind to the Ig-like 1 and Fz-like domains of MuSK, stabilizing it [29], and its absence results in decreased muscle membrane-bound MuSK [29, 36, 37]. However, this control of MuSK by ColQ has also been hypothesised to act via LRP4 [44]. In addition, the binding of ColQ to MuSK inhibits AGRIN signalling [29], demonstrating the opposing ways ColQ maintains NMJ homeostasis. This integral relationship between ColQ and MuSK led us to hypothesize that 3B2 treatment might stabilize and phosphorylate MuSK, thus providing a therapeutic benefit to *ColQ*^-/-^ mice. We found that 3B2 was unable to rescue many of the cellular and phenotypic characteristics of the *ColQ^-/-^* mice, including bodyweight, grip strength and fibre type switching, suggesting that this treatment may not be appropriate for *ColQ*-CMS. Because the absence of ColQ leads to reduced synaptic localization of MuSK [23, 37], *ColQ*^-/-^ mice had lower pMuSK levels in the TA lysate. However, without MuSK stabilization in the absence of ColQ, 3B2 treatment may not be able to increase pMuSK levels to provide therapeutic benefit in *ColQ*^-/-^ mice. Instead, 3B2 treatment increased total MuSK levels, but this did not improve the phenotypes of *ColQ*^-/-^ mice.

As previously reported [23, 36], *ColQ*^-/-^ mice had decreased bodyweight and motor strength, a lower proportion of Type I muscle fibres, and abnormal NMJ morphology. 3B2 treatment had no effect on bodyweight, motor strength, fibre size or type. In addition, it did not rescue any of the NMJ morphological changes in the *ColQ*^-/-^ (Mot) mice. In fact, many of the NMJ morphological parameters decreased further in *ColQ*^-/-^ (3B2) animals. While the absence of ColQ has been shown to reduce pMuSK levels *in vitro* [44], this is the first time it has been reported *in vivo.* However, at both the synapse and in the muscle overall, 3B2 increased MuSK levels, which was not accompanied by an increase in pMuSK.

The main mechanism of disease in *ColQ-*CMS is the absence of AChE, leading to an excess of ACh in the synaptic cleft, repetitive action potentials at the muscle fibre, and degeneration of the postjunctional folds. The stabilization of MuSK on the muscle membrane by ColQ may explain some of the common clinical features between *ColQ-*CMS and CMS subtypes directly impacted by the AGRIN/LRP4/MuSK pathway, such as *Agrn-* and *Dok7-*CMS [9, 22, 24, 26]. Furthermore, the most beneficial treatment for *ColQ-*CMS and CMS subtypes directly impacted by the AGRIN/LRP4/MuSK pathway is salbutamol, a treatment that helps to increase AChR clusters [10, 23, 48], although its mechanism of action is unknown. Unfortunately, 3B2 treatment did not improve any of the cellular and phenotypic characteristics in *ColQ^-/-^* mice, likely due to the reduction in pMuSK not being the main mechanism of disease of *ColQ-*CMS. These divergent results in *Agrn^nmf380^* and *ColQ*^-/-^ mice are supported by the well-known fact that treatments for CMS are specific to the mutated gene, and administering the wrong treatment to a patient can be detrimental. Thus, it remains important that new CMS treatments be tested in a variety of CMS models before being given to patients. In this case, we would hypothesize that 3B2 may not be beneficial for *ColQ-*CMS patients but may be extremely beneficial to *Agrn*-CMS patients. It will be important to test this compound in animal models with other mutations: we would suggest other components that act in the AGRIN/MuSK/LRP4 pathway as a good place to start. Indeed, clinical trials with a similar MuSK agonist antibody [46] are being conducted for patients with *Dok7-*CMS (NCT06436742). Studies also suggest that MuSK agonist antibodies may have use beyond CMS including MuSK-myasthenia gravis, ALS, and SMA [6, 13, 20, 30, 34, 41].

In this study, we tested the hypothesis that 3B2, a MuSK agonist antibody and derivative of ARGX-119, would be beneficial in animal models of CMS with defects in proteins that directly and indirectly interact with the AGRIN/LRP4/MuSK pathway. While the different results in *Agrn^nmf380^* and *ColQ*^-/-^ mice likely stem from the distinct functions of Agrin and ColQ, the two models received different treatment protocols due to phenotype severity. *Agrn^nmf380^* mice began treatment before the phenotype was visually evident at P5, while treatment of the *ColQ*^-/-^ mice began after symptoms, notably their smaller size, was evident and at P22, after NMJs in WT mice appear mature. Therefore, we cannot exclude the possibility that earlier dosing of the *ColQ*^-/-^ mice may lead to improved outcomes. However, we previously rescued some CMS features in *ColQ*^-/-^ mice using salbutamol, initiating treatment at a similar age [23], demonstrating that NMJs remain tractable post-symptom onset. Another difference between the treatment protocols was the dosing regimen: *Agrn^nmf380^*mice were injected at P5, P15 and P35, whereas *ColQ*^-/-^ mice were injected weekly. While we think these differences do not impact the overall conclusions of this study, follow-up experiments should be performed to determine whether alternative dosing might improve the deficits of *ColQ*^-/-^ mice, and whether starting treatment later would still rescue *Agrn^nmf380^*mice.

### Conclusions

In conclusion, successful treatment of CMS-mice with a MuSK agonist antibody (3B2) is mutation-dependent. In *Agrn^nmf380^* mice, treatment with the MuSK agonist antibody rescued NMJ morphology, cellular disease, and life span, while similar benefits were not found in *ColQ*^-/-^ mice. These results highlight the importance of identifying the genetic cause of CMS to effectively treat patients.

## Declarations

### Data Availability

The raw data supporting the conclusions of this article is available from the corresponding author, upon reasonable request.

### Ethical Approval

All animal procedures were performed following the approval of the University of Ottawa Animal Care Committee (experimental protocol 3120) and complied with the guidelines of the Canadian Council on Animal Care and the Animals for Research Act.

## Author Contributions

Conceptualization, H. L., S. S., S.B., R.V.

Methodology, K.H., O.A., R.C., S.S., J.Z., R.R., D.O., L.D.C.,

Formal analysis, S.S., K.H., O.A.

Resources, H. L., S.S., R.V.

Data curation, L.D.C., S.S., K.H., O.A.

Writing—original draft preparation, S.S., K.H., O.A.

Review and editing H.L., S.V., S.B., R.C., K.H., O.A., J.V., R.R., L.D.C., D.O., R.V.

Final draft preparation and writing, S.S., K.H., H.L.,

Supervision, S.S. and H. L.

Project administration, S. S. and H. L

Funding acquisition, H. L. and S.S.

## Acknowledgements

We would like to thank Robert Burgess for supplying the *Agrn^nmf380^* mouse and the Krejci laboratory for the *ColQ^-/-^* mouse. The BA-F8, SC-71, and BF-F3monoclonal antibodies deposited by Schiaffino, S., and the 6H1 monoclonal antibody deposited by Lucas, C., were obtained from the Developmental Studies Hybridoma Bank, created by the NICHD of the NIH and maintained at The University of Iowa, Department of Biology, Iowa City, IA 52242.

## References

1. Beeson D, Higuchi O, Palace J, Cossins J, Spearman H, Maxwell S, Newsom-Davis J, Burke G, Fawcett P, Motomura M, Müller JS, Lochmüller H, Slater C, Vincent A, Yamanashi Y (2006) Dok-7 mutations underlie a neuromuscular junction synaptopathy. Science 313:1975–1978. 10.1126/science.1130837

2. Bergamin E, Hallock PT, Burden SJ, Hubbard SR (2010) The cytoplasmic adaptor protein Dok7 activates the receptor tyrosine kinase MuSK via dimerization. Mol Cell 39:100–109. 10.1016/j.molcel.2010.06.007

3. Bogdanik LP, Burgess RW (2011) A valid mouse model of AGRIN-associated congenital myasthenic syndrome. Hum Mol Genet 20:4617–4633. 10.1093/hmg/ddr396

4. Borges LS, Yechikhov S, Lee YI, Rudell JB, Friese MB, Burden SJ, Ferns MJ (2008) Identification of a motif in the acetylcholine receptor beta subunit whose phosphorylation regulates rapsyn association and postsynaptic receptor localization. J Neurosci 28:11468–11476. 10.1523/JNEUROSCI.2508-08.2008

5. Burden SJ, Yumoto N, Zhang W (2013) The role of MuSK in synapse formation and neuromuscular disease. Cold Spring Harb Perspect Biol 5:a009167. 10.1101/cshperspect.a009167

6. Cantor S, Zhang W, Delestrée N, Remédio L, Mentis GZ, Burden SJ (2018) Preserving neuromuscular synapses in ALS by stimulating MuSK with a therapeutic agonist antibody. Elife 7:e34375. 10.7554/eLife.34375

7. Cartaud A, Strochlic L, Guerra M, Blanchard B, Lambergeon M, Krejci E, Cartaud J, Legay C (2004) MuSK is required for anchoring acetylcholinesterase at the neuromuscular junction. J Cell Biol 165:505–515. 10.1083/jcb.200307164

8. Chen B-H, Lin Z-Y, Zeng X-X, Jiang Y-H, Geng F (2024) LRP4-related signalling pathways and their regulatory role in neurological diseases. Brain Res 1825:148705. 10.1016/j.brainres.2023.148705

9. Chevessier F, Girard E, Molgó J, Bartling S, Koenig J, Hantaï D, Witzemann V (2008) A mouse model for congenital myasthenic syndrome due to MuSK mutations reveals defects in structure and function of neuromuscular junctions. Hum Mol Genet 17:3577–3595. 10.1093/hmg/ddn251

10. Clausen L, Cossins J, Beeson D (2018) Beta-2 Adrenergic Receptor Agonists Enhance AChR Clustering in C2C12 Myotubes: Implications for Therapy of Myasthenic Disorders. J Neuromuscul Dis 5:231–240. 10.3233/JND-170293

11. Donger C, Krejci E, Serradell AP, Eymard B, Bon S, Nicole S, Chateau D, Gary F, Fardeau M, Massoulié J, Guicheney P (1998) Mutation in the human acetylcholinesterase-associated collagen gene, COLQ, is responsible for congenital myasthenic syndrome with end-plate acetylcholinesterase deficiency (Type Ic). Am J Hum Genet 63:967–975. 10.1086/302059

12. Feng G, Krejci E, Molgo J, Cunningham JM, Massoulié J, Sanes JR (1999) Genetic Analysis of Collagen Q: Roles in Acetylcholinesterase and Butyrylcholinesterase Assembly and in Synaptic Structure and Function. The Journal of Cell Biology 144:1349–1360. 10.1083/jcb.144.6.1349

13. Feng Z, Lam S, Tenn E-MS, Ghosh AS, Cantor S, Zhang W, Yen P-F, Chen KS, Burden S, Paushkin S, Ayalon G, Ko C-P (2021) Activation of Muscle-Specific Kinase (MuSK) Reduces Neuromuscular Defects in the Delta7 Mouse Model of Spinal Muscular Atrophy (SMA). Int J Mol Sci 22:8015. 10.3390/ijms22158015

14. Friese MB, Blagden CS, Burden SJ (2007) Synaptic differentiation is defective in mice lacking acetylcholine receptor beta-subunit tyrosine phosphorylation. Development 134:4167–4176. 10.1242/dev.010702

15. Hallock PT, Xu C-F, Park T-J, Neubert TA, Curran T, Burden SJ (2010) Dok-7 regulates neuromuscular synapse formation by recruiting Crk and Crk-L. Genes Dev 24:2451–2461. 10.1101/gad.1977710

16. Herbst R, Huijbers MG, Oury J, Burden SJ (2024) Building, Breaking, and Repairing Neuromuscular Synapses. Cold Spring Harb Perspect Biol 16:a041490. 10.1101/cshperspect.a041490

17. Jacquier A, Risson V, Simonet T, Roussange F, Lacoste N, Ribault S, Carras J, Theuriet J, Girard E, Grosjean I, Le Goff L, Kröger S, Meltoranta J, Bauché S, Sternberg D, Fournier E, Kostera-Pruszczyk A, O’Connor E, Eymard B, Lochmüller H, Martinat C, Schaeffer L (2022) Severe congenital myasthenic syndromes caused by agrin mutations affecting secretion by motoneurons. Acta Neuropathol 144:707–731. 10.1007/s00401-022-02475-8

18. Jones RA, Reich CD, Dissanayake KN, Kristmundsdottir F, Findlater GS, Ribchester RR, Simmen MW, Gillingwater TH (2016) NMJ-morph reveals principal components of synaptic morphology influencing structure-function relationships at the neuromuscular junction. Open Biol 6:160240. 10.1098/rsob.160240

19. Kim N, Burden SJ (2008) MuSK controls where motor axons grow and form synapses. Nat Neurosci 11:19–27. 10.1038/nn2026

20. Lim JL, Jensen SM, Plomp JJ, Vankerckhoven B, Kneip C, Coppejans R, Steyaert C, Moens K, De Clercq L, Tannemaat MR, Ulrichts P, Silence K, van der Maarel SM, Vergoossen DLE, Vanhauwaert R, Verschuuren JJ, Huijbers MG (2025) Patient-specific therapeutic benefit of MuSK agonist antibody ARGX-119 in MuSK myasthenia gravis passive transfer models. iScience 28:111684. 10.1016/j.isci.2024.111684

21. Lin S, Maj M, Bezakova G, Magyar JP, Brenner HR, Ruegg MA (2008) Muscle-wide secretion of a miniaturized form of neural agrin rescues focal neuromuscular innervation in agrin mutant mice. Proc Natl Acad Sci U S A 105:11406–11411. 10.1073/pnas.0801683105

22. Maselli RA, Arredondo J, Cagney O, Ng JJ, Anderson JA, Williams C, Gerke BJ, Soliven B, Wollmann RL (2010) Mutations in MUSK causing congenital myasthenic syndrome impair MuSK-Dok-7 interaction. Hum Mol Genet 19:2370–2379. 10.1093/hmg/ddq110

23. McMacken GM, Spendiff S, Whittaker RG, O’Connor E, Howarth RM, Boczonadi V, Horvath R, Slater CR, Lochmüller H (2019) Salbutamol modifies the neuromuscular junction in a mouse model of ColQ myasthenic syndrome. Hum Mol Genet 28:2339–2351. 10.1093/hmg/ddz059

24. Mihaylova V, Müller JS, Vilchez JJ, Salih MA, Kabiraj MM, D’Amico A, Bertini E, Wölfle J, Schreiner F, Kurlemann G, Rasic VM, Siskova D, Colomer J, Herczegfalvi A, Fabriciova K, Weschke B, Scola R, Hoellen F, Schara U, Abicht A, Lochmüller H (2008) Clinical and molecular genetic findings in COLQ-mutant congenital myasthenic syndromes. Brain 131:747–759. 10.1093/brain/awm325

25. Müller JS, Herczegfalvi A, Vilchez JJ, Colomer J, Bachinski LL, Mihaylova V, Santos M, Schara U, Deschauer M, Shevell M, Poulin C, Dias A, Soudo A, Hietala M, Aärimaa T, Krahe R, Karcagi V, Huebner A, Beeson D, Abicht A, Lochmüller H (2007) Phenotypical spectrum of DOK7 mutations in congenital myasthenic syndromes. Brain 130:1497–1506. 10.1093/brain/awm068

26. Ohkawara B, Cabrera-Serrano M, Nakata T, Milone M, Asai N, Ito K, Ito M, Masuda A, Ito Y, Engel AG, Ohno K (2014) LRP4 third β-propeller domain mutations cause novel congenital myasthenia by compromising agrin-mediated MuSK signaling in a position-specific manner. Hum Mol Genet 23:1856–1868. 10.1093/hmg/ddt578

27. Ohno K, Brengman J, Tsujino A, Engel AG (1998) Human endplate acetylcholinesterase deficiency caused by mutations in the collagen-like tail subunit (ColQ) of the asymmetric enzyme. Proc Natl Acad Sci U S A 95:9654–9659. 10.1073/pnas.95.16.9654

28. Ohno K, Ohkawara B, Shen X-M, Selcen D, Engel AG (2023) Clinical and Pathologic Features of Congenital Myasthenic Syndromes Caused by 35 Genes-A Comprehensive Review. Int J Mol Sci 24:3730. 10.3390/ijms24043730

29. Otsuka K, Ito M, Ohkawara B, Masuda A, Kawakami Y, Sahashi K, Nishida H, Mabuchi N, Takano A, Engel AG, Ohno K (2015) Collagen Q and anti-MuSK autoantibody competitively suppress agrin/LRP4/MuSK signaling. Sci Rep 5:13928. 10.1038/srep13928

30. Oury J, Gamallo-Lana B, Santana L, Steyaert C, Vergoossen DLE, Mar AC, Vankerckhoven B, Silence K, Vanhauwaert R, Huijbers MG, Burden SJ (2024) Agonist antibody to MuSK protects mice from MuSK myasthenia gravis. Proc Natl Acad Sci U S A 121:e2408324121. 10.1073/pnas.2408324121

31. Oury J, Zhang W, Leloup N, Koide A, Corrado AD, Ketavarapu G, Hattori T, Koide S, Burden SJ (2021) Mechanism of disease and therapeutic rescue of Dok7 congenital myasthenia. Nature 595:404–408. 10.1038/s41586-021-03672-3

32. Sander A, Hesser BA, Witzemann V (2001) MuSK induces in vivo acetylcholine receptor clusters in a ligand-independent manner. J Cell Biol 155:1287–1296. 10.1083/jcb.200105034

33. Schiaffino S, Reggiani C (2011) Fiber types in mammalian skeletal muscles. Physiol Rev 91:1447–1531. 10.1152/physrev.00031.2010

34. Sengupta-Ghosh A, Dominguez SL, Xie L, Barck KH, Jiang Z, Earr T, Imperio J, Phu L, Budayeva HG, Kirkpatrick DS, Cai H, Eastham-Anderson J, Ngu H, Foreman O, Hedehus M, Reichelt M, Hotzel I, Shang Y, Carano RAD, Ayalon G, Easton A (2019) Muscle specific kinase (MuSK) activation preserves neuromuscular junctions in the diaphragm but is not sufficient to provide a functional benefit in the SOD1G93A mouse model of ALS. Neurobiol Dis 124:340–352. 10.1016/j.nbd.2018.12.002

35. Shen C, Lu Y, Zhang B, Figueiredo D, Bean J, Jung J, Wu H, Barik A, Yin D-M, Xiong W-C, Mei L (2013) Antibodies against low-density lipoprotein receptor-related protein 4 induce myasthenia gravis. J Clin Invest 123:5190–5202. 10.1172/JCI66039

36. Sigoillot SM, Bourgeois F, Karmouch J, Molgó J, Dobbertin A, Chevalier C, Houlgatte R, Léger J, Legay C (2016) Neuromuscular junction immaturity and muscle atrophy are hallmarks of the ColQ-deficient mouse, a model of congenital myasthenic syndrome with acetylcholinesterase deficiency. FASEB J 30:2382–2399. 10.1096/fj.201500162

37. Sigoillot SM, Bourgeois F, Lambergeon M, Strochlic L, Legay C (2010) ColQ controls postsynaptic differentiation at the neuromuscular junction. J Neurosci 30:13–23. 10.1523/JNEUROSCI.4374-09.2010

38. Slater CR (2008) Reliability of neuromuscular transmission and how it is maintained. Handb Clin Neurol 91:27–101. 10.1016/S0072-9752(07)01502-3

39. Slater CR (2020) “Fragmentation” of NMJs: a sign of degeneration or regeneration? A long journey with many junctions. Neuroscience 439:28–40. 10.1016/j.neuroscience.2019.05.017

40. Spendiff S, Howarth R, McMacken G, Davey T, Quinlan K, O’Connor E, Slater C, Hettwer S, Mäder A, Roos A, Horvath R, Lochmüller H (2020) Modulation of the Acetylcholine Receptor Clustering Pathway Improves Neuromuscular Junction Structure and Muscle Strength in a Mouse Model of Congenital Myasthenic Syndrome. Front Mol Neurosci 13:594220. 10.3389/fnmol.2020.594220

41. Sun S, Shen Y, Zhang X, Ding N, Xu Z, Zhang Q, Li L (2024) The MuSK agonist antibody protects the neuromuscular junction and extends the lifespan in C9orf72-ALS mice. Mol Ther 32:2176–2189. 10.1016/j.ymthe.2024.05.016

42. Theuriet J, Masingue M, Behin A, Ferreiro A, Bassez G, Jaubert P, Tarabay O, Fer F, Pegat A, Bouhour F, Svahn J, Petiot P, Jomir L, Chauplannaz G, Cornut-Chauvinc C, Manel V, Salort-Campana E, Attarian S, Fortanier E, Verschueren A, Kouton L, Camdessanché J-P, Tard C, Magot A, Péréon Y, Noury J-B, Minot-Myhie M-C, Perie M, Taithe F, Farhat Y, Millet A-L, Cintas P, Solé G, Spinazzi M, Esselin F, Renard D, Sacconi S, Ezaru A, Malfatti E, Mallaret M, Magy L, Diab E, Merle P, Michaud M, Fournier M, Pakleza AN, Chanson J-B, Lefeuvre C, Laforet P, Richard P, Sternberg D, Villar-Quiles R-N, Stojkovic T, Eymard B (2024) Congenital myasthenic syndromes in adults: clinical features, diagnosis and long-term prognosis. Brain 147:3849–3862. 10.1093/brain/awae124

43. Thompson R, Bonne G, Missier P, Lochmüller H (2019) Targeted therapies for congenital myasthenic syndromes: systematic review and steps towards a treatabolome. Emerg Top Life Sci 3:19–37. 10.1042/ETLS20180100

44. Uyen Dao TM, Barbeau S, Messéant J, Della-Gaspera B, Bouceba T, Semprez F, Legay C, Dobbertin A (2023) The collagen ColQ binds to LRP4 and regulates the activation of the Muscle-Specific Kinase-LRP4 receptor complex by agrin at the neuromuscular junction. J Biol Chem 299:104962. 10.1016/j.jbc.2023.104962

45. Valenzuela DM, Stitt TN, DiStefano PS, Rojas E, Mattsson K, Compton DL, Nuñez L, Park JS, Stark JL, Gies DR (1995) Receptor tyrosine kinase specific for the skeletal muscle lineage: expression in embryonic muscle, at the neuromuscular junction, and after injury. Neuron 15:573–584. 10.1016/0896-6273(95)90146-9

46. Vanhauwaert R, Oury J, Vankerckhoven B, Steyaert C, Jensen SM, Vergoossen DLE, Kneip C, Santana L, Lim JL, Plomp JJ, Augustinus R, Koide S, Blanchetot C, Ulrichts P, Huijbers MG, Silence K, Burden SJ (2024) ARGX-119 is an agonist antibody for human MuSK that reverses disease relapse in a mouse model of congenital myasthenic syndrome. Sci Transl Med 16:eado7189. 10.1126/scitranslmed.ado7189

47. Viegas S, Jacobson L, Waters P, Cossins J, Jacob S, Leite MI, Webster R, Vincent A (2012) Passive and active immunization models of MuSK-Ab positive myasthenia: electrophysiological evidence for pre and postsynaptic defects. Exp Neurol 234:506–512. 10.1016/j.expneurol.2012.01.025

48. Webster RG, Vanhaesebrouck AE, Maxwell SE, Cossins JA, Liu W, Ueta R, Yamanashi Y, Beeson DMW (2020) Effect of salbutamol on neuromuscular junction function and structure in a mouse model of DOK7 congenital myasthenia. Hum Mol Genet 29:2325–2336. 10.1093/hmg/ddaa116

49. Willadt S, Nash M, Slater CR (2016) Age-related fragmentation of the motor endplate is not associated with impaired neuromuscular transmission in the mouse diaphragm. Sci Rep 6:24849. 10.1038/srep24849

50. Willmann R, Dubach J, Chen K (2011) Developing standard procedures for pre-clinical efficacy studies in mouse models of spinal muscular atrophy. Neuromuscular Disorders 21:74–77. 10.1016/j.nmd.2010.09.014

51. Yumoto N, Kim N, Burden SJ (2012) Lrp4 is a retrograde signal for presynaptic differentiation at neuromuscular synapses. Nature 489:438–442. 10.1038/nature11348

52. Zhao Y, Li Y, Bian Y, Yao S, Liu P, Yu M, Zhang W, Wang Z, Yuan Y (2021) Congenital myasthenic syndrome in China: genetic and myopathological characterization. Ann Clin Transl Neurol 8:898–907. 10.1002/acn3.51346

53. Zong Y, Jin R (2013) Structural mechanisms of the agrin-LRP4-MuSK signaling pathway in neuromuscular junction differentiation. Cell Mol Life Sci 70:3077–3088. 10.1007/s00018-012-1209-9

54. Mira Vision. https://www.mira.vision/. Accessed 29 Jun 2023

