## Supplemental Material for "Gene-specific response to MuSK agonist antibody in the treatment of Congenital Myasthenic Syndromes"

**Supplementary material**

**Supplementary figures**

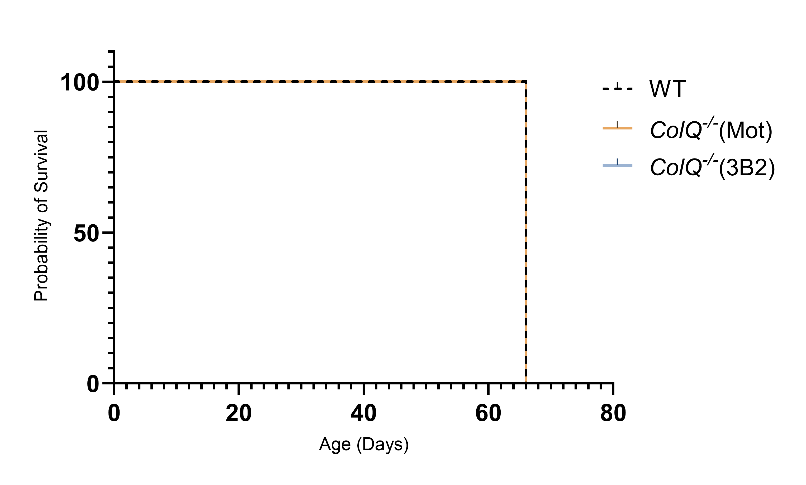

**Supplementary Figure 1. Survival of *ColQ^-/-^* mice.** All animals survived to the end of the study at P66. WT n=10, *ColQ*^-/-^ (Mot) n=10, *ColQ*^-/-^ (3B2) n=12

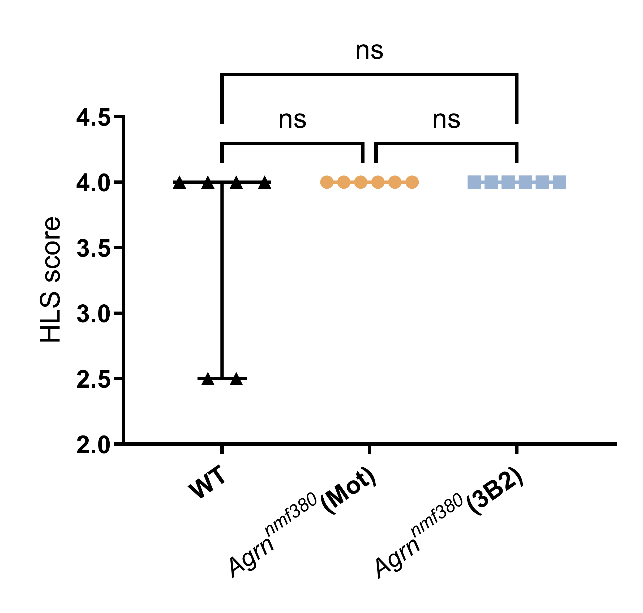

**Supplementary Figure 2. Hindlimb suspension score (HLS) in *Agrn^nmf380^* and WT mice.** There were no differences in the HLS score between groups. Median and IQR, Kruskal-Wallis test with Dunn’s correction for multiple comparisons. WT n=6, *Agrn^nmf380^* (Mot) n=6, *Agrn^nmf380^* (3B2) n=6.

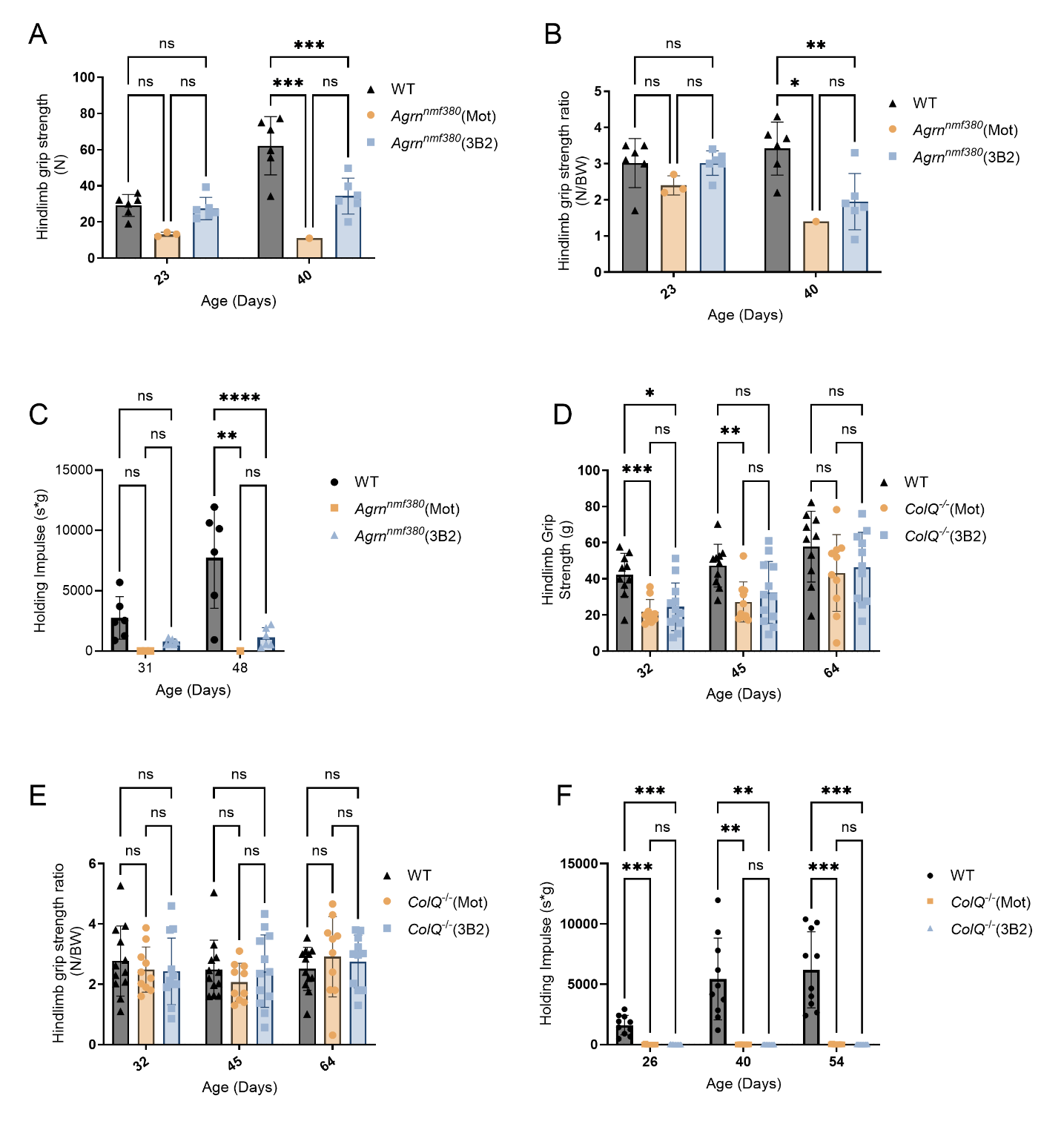

**Supplementary Figure 3. Strength assessments in *Agrn^nmf380^* and *ColQ^-/-^* animals (A)** Hindlimb grip strength revealed differences between WT and *Agrn^nmf380^* mice at P40. **(B)** There were no differences in hindlimb grip strength ratio between *Agrn^nmf380^* animals treated with Mot or 3B2. **(C)** To determine the holding impulse, the time to fall in seconds was multipled by the bodyweight of the animal. WT animals were not significantly stronger than *Agrn^nmf380^* (Mot) or *Agrn^nmf380^* (3B2) animals until P48. Although the average holding ratio in *Agrn^nmf380^* mice went from 0 to 3.5 with 3B2 treatment, there was no significant difference. **(D)** WT animals had greater hindlimb strength than *ColQ^-/-^* animals, though these differences disappeared when normalized for body weight (**E**) and no improvements were seen with 3B2 treatment. **(F)** A similar pattern was observed in holding impulse. 2-Way ANOVA with Tukey’s multiple comparisons correction. At P23 & 31 WT n=6, *Agrn^nmf380^* (Mot) n=3, *Agrn^nmf380^* (3B2) n=6, at P42 &P48 WT n=6, *Agrn^nmf380^* (Mot) n=1, *Agrn^nmf380^* (3B2) n=6. WT n=10, *ColQ*^-/-^ (Mot) n=10, *ColQ*^-/-^ (3B2) n=12. Graphs show means and error bars indicate sd.

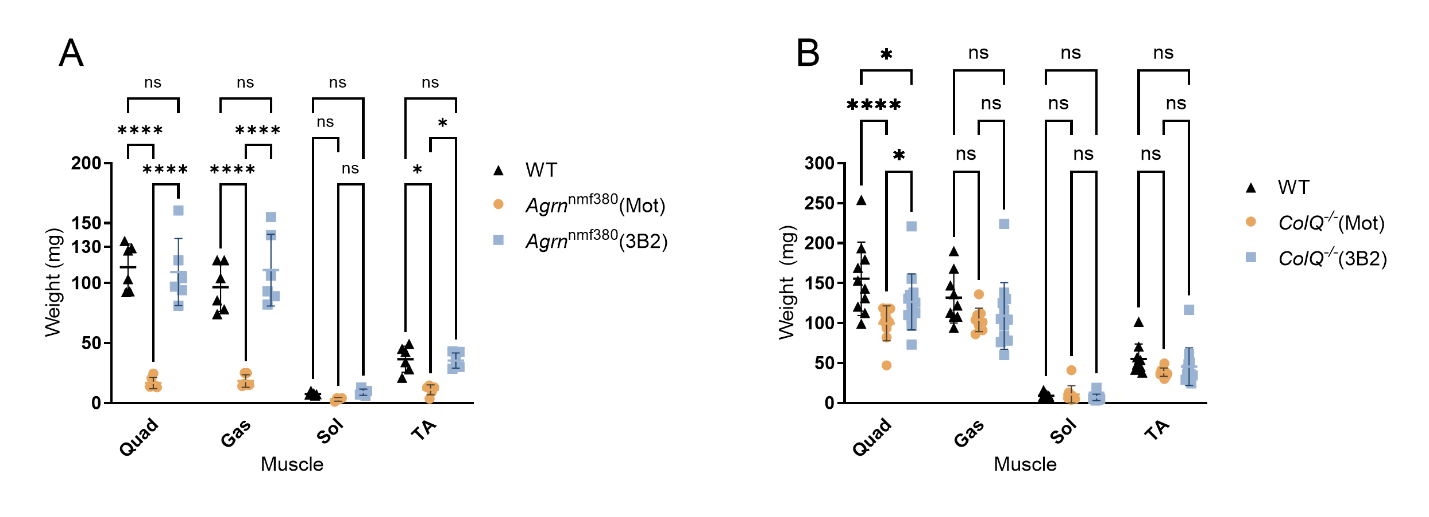
 **Supplementary Figure 4. Raw values muscle weight in WT, 3B2 treated and untreated animals.** **(A)** 3B2 treatment fully rescued quad, gas, and TA muscle weights of *Agrn^nmf380^* mice back to WT levels. WT n=6, *Agrn^nmf380^* (Mot) n=6, *Agrn^nmf380^* (3B2) n=6. **(B)** *ColQ^-/-^* (3B2) mice had heavier quad muscles compared to *ColQ^-/-^* (Mot) mice. WT n=10, *ColQ*^-/-^ (Mot) n=10, *ColQ*^-/-^ (3B2) n=12. 2-Way ANOVA with Tukey’s multiple comparisons correction. Graphs show means and error bars indicate sd. *p<0.05, **p<0.005, ***p<0.001, ****p<0.0001.

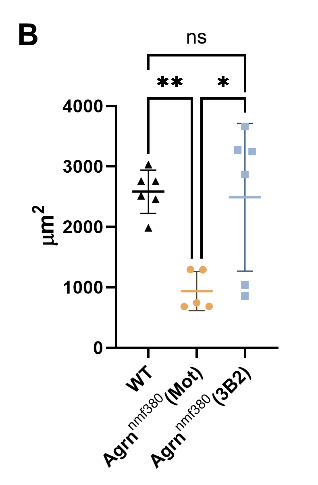

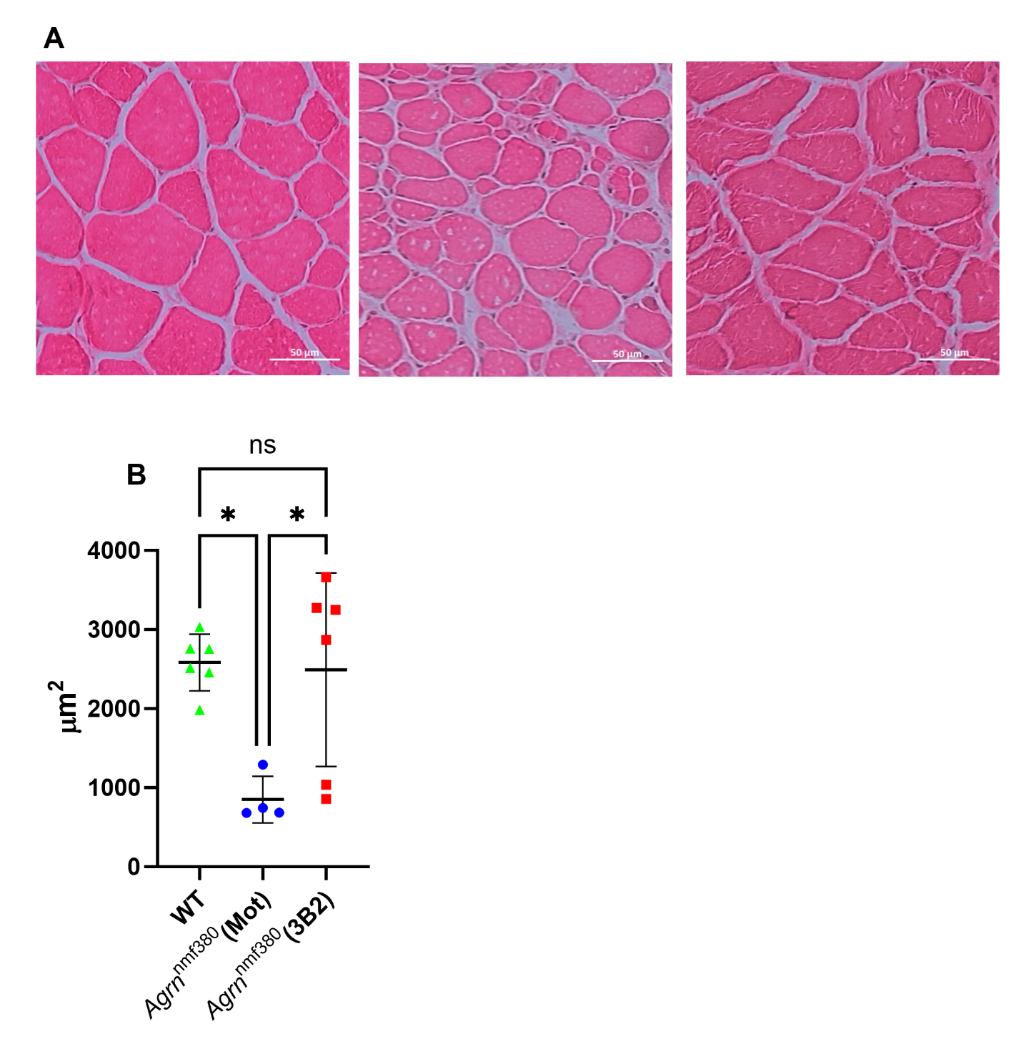

**Supplementary Figure 5. Quadriceps fibre size in *Agrn^nmf380^* mice. (A)** Haematoxylin and eosin was used to label muscle cross sections from WT (left), *Agrn^nmf380^* (Mot) (middle), and *Agrn^nmf380^* (3B2) (right) mice quadriceps. **(B)** *Agrn^nmf380^* (Mot) mice had significantly decreased myofiber size compared to both WT mice and *Agrn^nmf380^* (3B2) mice. ANOVA with Tukey’s multiple comparisons correction. Graphs show means and error bars indicate sd. WT n=6, *Agrn^nmf380^* (Mot) n=4, *Agrn^nmf380^* (3B2) n=6. *p<0.05

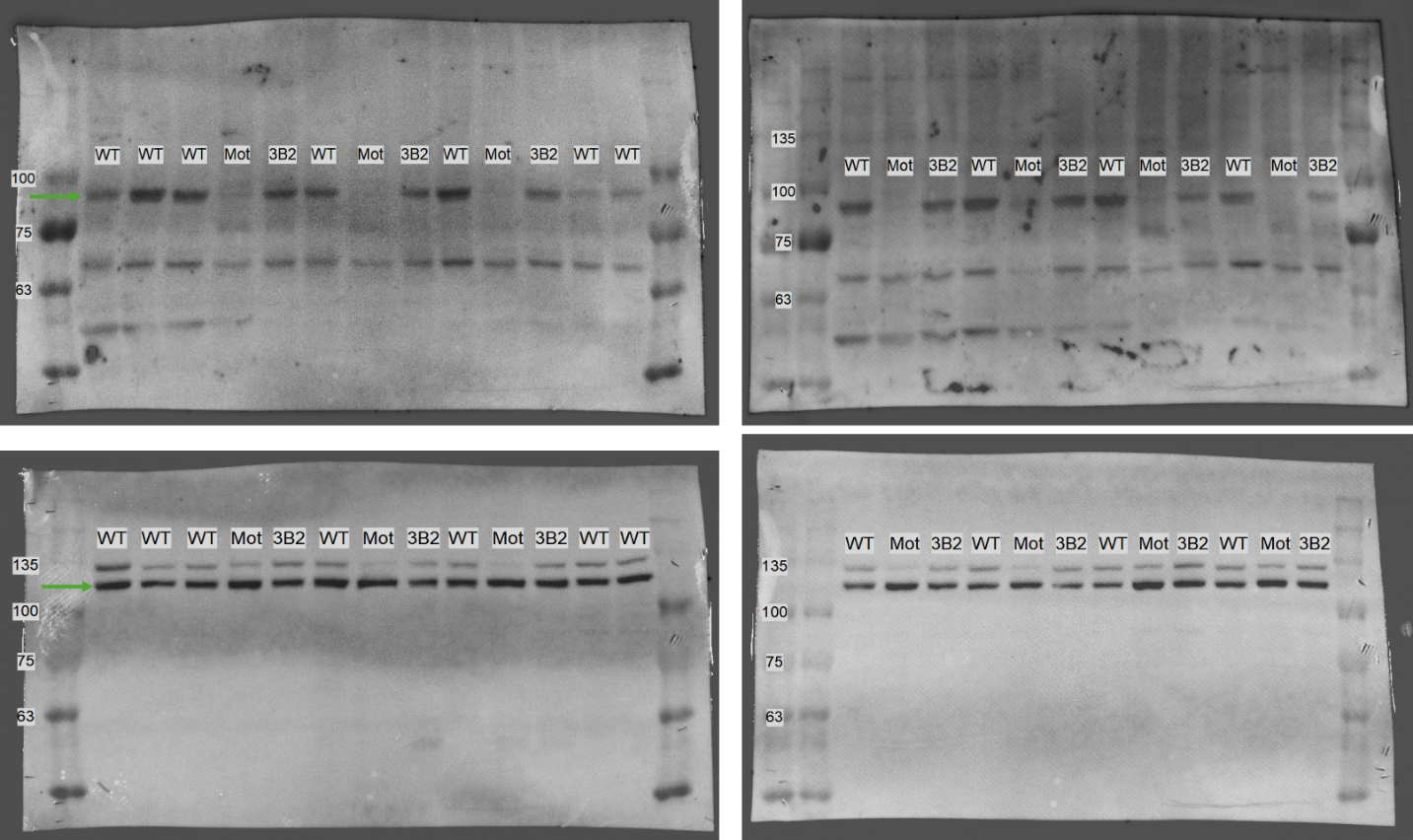

**Supplementary Figure 6. Full Western Blot images used to quantify pMuSK in WT and *Agrn^nmf380^* mice.** Bands were quantified and normalised to vinculin loading control. Some samples were run on more than one blot so that samples could be normalised across different blots. Thick ladder band is 75kDa and pMuSK is identified (top gels, green arrow) at 93kD. Loading control is vinculin (bottom gels, green arrow) at 120kD.

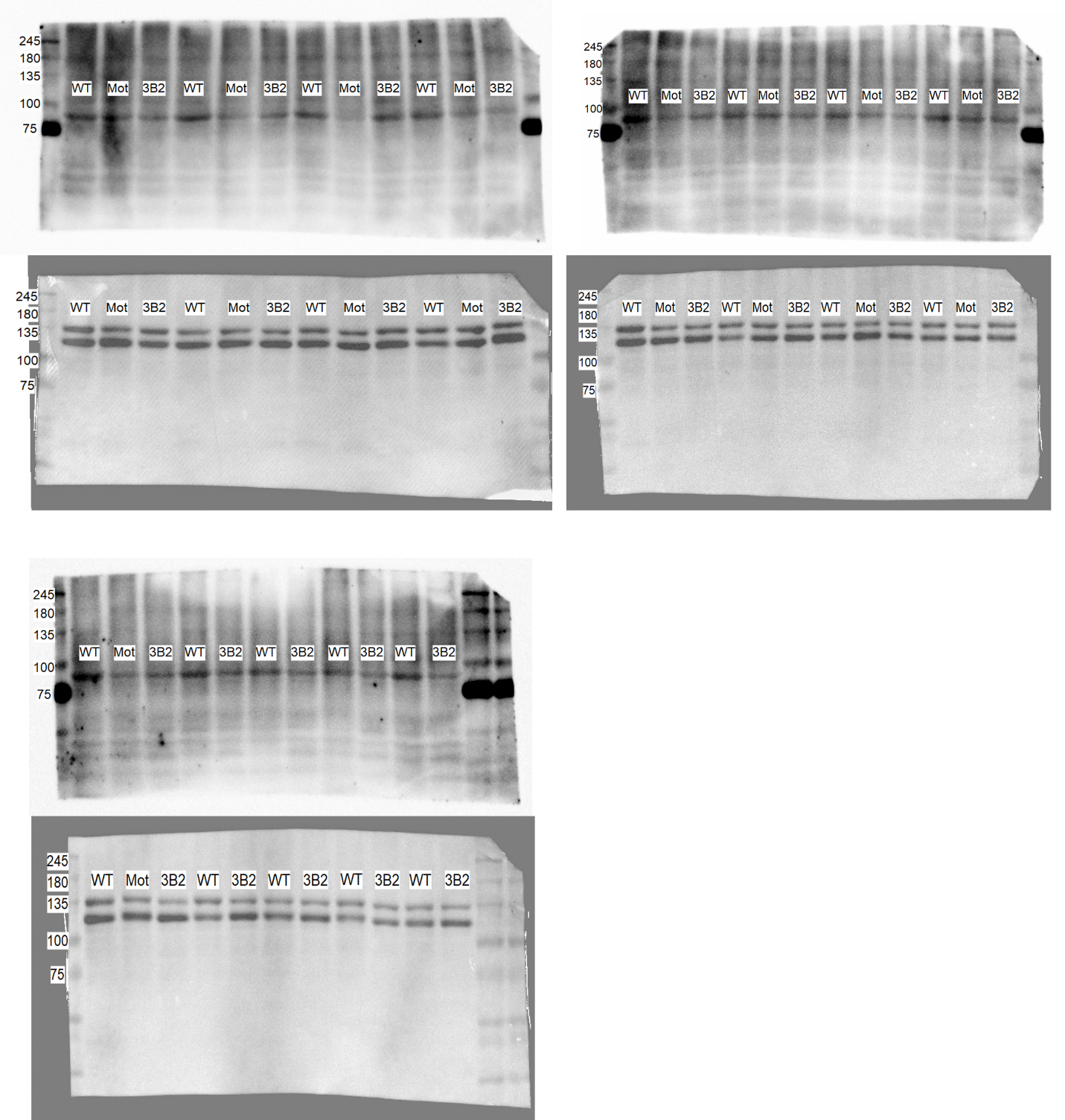
 **Supplementary Figure 7. Full Western blot images used to quantify pMuSK in WT and *ColQ^-/-^* mice.** Bands were quantified and normalised to vinculin loading control. Some samples were run on more than one blot so that samples could be normalised across different blots. Thick ladder band is 75KDa and the pMuSK band is visible at 93Kda (Top gels). Vinculin (bottom gels) was identified at 120kD.

**Supplementary tables**

**Supplementary Table 1: Treatment protocol showing timings and dosages for experimental, control, and WT mice.**

| **Model** | **N** | **Genotype** | **Treatment** | **Dose Level (mg/kg)** | **Route** | **Regimen** |
| --- | --- | --- | --- | --- | --- | --- |
| AGRN | 6 | *Agrn^nmf380^* | 3B2 | 20 mg/kg at P5, 10mg/kg at P15 and P35 | IP | P5, P15, P35 |
|  | 6 |  | Mota |  |  |  |
|  | 6 | Wild type | None | N/A | N/A | N/A |
| ColQ | 12 | *ColQ*^-/-^ mice | 3B2 | 20 mg/kg at P22, followed by weekly 10 mg/kg | IP | P22, P29, P36, P43, P50 and P57 |
|  | 10 |  | Mota |  |  |  |
|  | 10 | Wild type | None | N/A | N/A | N/A |

**Supplementary Table 2: Serum concentrations of 3B2 in WT and *Agrn^nmf380^* mice**.

| **Group** | **Animal ID** | **Dosing** | **Sample point** | **[3B2] (µg/mL)** | **Sample point** | **[3B2] (****µg/mL)** |
| --- | --- | --- | --- | --- | --- | --- |
| WT | T2M1 | NA | P35 | <LLOQ | P50 | <LLOQ |
|  | T2F2 |  |  | NS |  | <LLOQ |
|  | Y1M2 |  |  | NS |  | <LLOQ |
|  | Y1M3 |  |  | NS |  | <LLOQ |
|  | Y1M4 |  |  | NS |  | <LLOQ |
|  | Y1M6 |  |  | NS |  | <LLOQ |
|  | Y1F1 |  |  | NS |  | <LLOQ |
|  | AK3F1 |  |  | <LLOQ |  | <LLOQ |
|  | AK3M2 |  |  | <LLOQ |  | <LLOQ |
|  | AK3M3 |  |  | <LLOQ |  | <LLOQ |
| *Agrn^nmf380^* (Mot) | X1M3 | 20 mg/kg at P5, 10mg/kg at P15 and P35 | P35 | NS | P50 | <LLOQ |
|  | AH3F3 |  |  | NS |  | NS |
|  | AH3F5 |  |  | NS |  | NS |
|  | AJ2F1 |  |  | NS |  | <LLOQ |
|  | AJ5M2 |  |  | NS |  | <LLOQ |
|  | AJ5M4 |  |  | NS |  | <LLOQ |
| *Agrn^nmf380^* (3B2) | AG3F1 | 20 mg/kg at P5, 10mg/kg at P15 and P35 | P35 | 12.9 | P50 | 44.05 |
|  | AJ5F3 |  |  | 3.91 |  | NS |
|  | AJ5F5 |  |  | 7.79 |  | 4.00 |
|  | AL2M4 |  |  | 10.27 |  | 15.20 |
|  | AG3M2 |  |  | 12.54 |  | 36.80 |
|  | AG3M4 |  |  | 1.94 |  | 39.74 |

ELISAs were performed on serum to determine the levels of 3B2. As expected, levels of 3B2 in the WT and *Agrn^nmf380^* (Mot) mice were undetectable, but detectable in the *Agrn^nmf380^* (3B2) animals at both P35 and P50. Furthermore, there was a significant increase in 3B2 serum levels in the *Agrn^nmf380^* (3B2) mice from P35 to P50, demonstrating the accumulation of the compound in the blood. **NS** no samples taken (Animal care guidelines dictate a minimum body weight that must be reached before blood collection can be performed), **LLOQ** Lower limit of quantification (3B2 0.1µg/mL), **NA** not applicable.

**Supplementary Table 3: Serum concentrations of 3B2 in WT and *ColQ^-/-^* mice.**

| **Group** | **Animal ID** | **Dosing** | **Sample point** | **[3B2] (µg/mL)** | **Sample point** | **[3B2] (µg/mL)** | **Sample point** | **[3B2](µg/mL)** |
| --- | --- | --- | --- | --- | --- | --- | --- | --- |
| WT | AW2F3 | NA | P29 | NS | P43 | NES | P66 | <LLOQ |
|  | AW4M1 |  |  | NS |  | <LLOQ |  | <LLOQ |
|  | BF2F3 |  |  | NS |  | NS |  | <LLOQ |
|  | BJ3F3 |  |  | NS |  | <LLOQ |  | <LLOQ |
|  | BJ3M2 |  |  | NS |  | NS |  | <LLOQ |
|  | BK1M4 |  |  | NS |  | <LLOQ |  | <LLOQ |
|  | BK2F1 |  |  | NS |  | <LLOQ |  | <LLOQ |
|  | BK2M4 |  |  | NS |  | NS |  | <LLOQ |
|  | BM2F1 |  |  | <LLOQ |  | <LLOQ |  | <LLOQ |
|  | BM2F2 |  |  | NS |  | <LLOQ |  | <LLOQ |
|  | BM2M1 |  |  | <LLOQ |  | <LLOQ |  | <LLOQ |
| *ColQ^-/-^* Mota | AW2M1 | 20 mg/kg at P22  followed by weekly 10 mg/kg | P29 | NS | P43 | NS | P66 | <LLOQ |
|  | AW2M2 |  |  | NS |  | NS |  | <LLOQ |
|  | BF2F1 |  |  | NS |  | NS |  | <LLOQ |
|  | BF2F2 |  |  | NS |  | <LLOQ |  | <LLOQ |
|  | BF2M3 |  |  | NS |  | NS |  | <LLOQ |
|  | BJ3F4 |  |  | NS |  | NS |  | <LLOQ |
|  | BK1M2 |  |  | NS |  | NS |  | <LLOQ |
|  | BK1M3 |  |  | NS |  | NS |  | <LLOQ |
|  | BK2F2 |  |  | NS |  | NS |  | <LLOQ |
|  | BX4F3 |  |  | NS |  | NS |  | <LLOQ |
| *ColQ^-/-^* 3B2 | AV2F1 | 20 mg/kg at P22  followed by weekly 10 mg/kg | P29 | NS | P43 | NS | P66 | 83.8 |
|  | AW4F4 |  |  | NS |  | NS |  | 15.3 |
|  | AW4M2 |  |  | NS |  | NS |  | 39.3 |
|  | AW4M4 |  |  | NS |  | 12.5 |  | 32.1 |
|  | BI1F2 |  |  | NS |  | <LLOQ |  | 88.4 |
|  | BJ1F3 |  |  | NS |  | NS |  | 91.9 |
|  | BJ1M4 |  |  | NS |  | NS |  | 70.8 |
|  | BJ2F3 |  |  | NS |  | NS |  | 86.5 |
|  | BJ4F2 |  |  | NS |  | NS |  | 94.8 |
|  | BJ4M2 |  |  | NS |  | 37.4 |  | 108.6 |
|  | BK2M2 |  |  | NS |  | NS |  | 96.7 |
|  | BK3M6 |  |  | NS |  | NS |  | 128.6 |
|  | BX2M1 |  |  | NS |  | NS |  | 138.6 |

Levels of circulating 3B2 were below the levels of detection in the WT and *ColQ*^-/-^ (Mot) animals. While circulating 3B2 was clearly detectible in the *ColQ*^-/-^ (3B2) animals, due to the lack of serum available at P43, we were unable to tell if there had been an accumulation. However, the two animals for which we had adequate samples at both P43 and P66, showed a more than 100% increase in circulating 3B2. The levels observed at the end of the study in the *ColQ*^-/-^ mice were considerably higher than those in the *Agrn^nmf380^* mice, suggesting the lack of changes observed in the *ColQ*^-/-^ (3B2) animals was not attributable to underdosing. **NS** no samples taken (Animal care guidelines dictate a minimum body weight that must be reached before blood collection can be performed), **LLOQ** Lower limit of quantification (3B2 0.1µg/mL), **NA** not applicable. **NES** not enough sample.

**Supplementary Table 4: Post hoc analysis of inverted hanging wire test.**

|  | **Non-holder** | **Holder** |
| --- | --- | --- |
| **WT** | 0 | 6 |
| ***Agrn^nmf380^* (Mot)** | 3 | 0 |
| ***Agrn^nmf380^*(3B2)** | 0 | 6 |

Initial analysis of inverted hanging wire test showed no significant differences in motor performance between *Agrn^nmf380^* (Mot) mice and *Agrn^nmf380^* (3B2) mice. Mice were classified as non-holders (<2 seconds) and holders (> 2 seconds). Fisher’s exact test was performed and showed there was a significant improvement in treated animals at P31, p=0.0022. n=6.

**Supplementary Table 5: Morphological variables of NMJs from *Agrn^nmf380^* mice**.

| **Variable** | **WT** | ***Agrn^nmf380^* Mot** | ***Agrn^nmf380^* 3B2** |
| --- | --- | --- | --- |
| **Core Variables** |  |  |  |
| **Presynaptic** |  |  |  |
| Absent Synaptophysin staining (%) | 0 | 0 | 0 |
| Nerve terminal area (µm^2^) | 184.3±69.5(71) | 105.3±92.9(28) ^**^ | 167.1±110.8(48) ^#^ |
| Nerve terminal perimeter (µm) | 216.6±65.8(72) | 147±105.2(28) ^*^ | 282.8±176.2(49) ^####^ |
| Number of terminal branches | 27±13.3(72) | 19.5±9.8(28) | 49.0±63(49) ^##^ |
| Number of branch points | 15.8±7.8(72) | 8.4±6.4(28) ^**^ | 9.6±8.1(49) ^*^ |
| Total length branches (µm) | 104.3±33.1(72) | 67.0±48.5(28) ^*^ | 114.2±73.1(49) ^###^ |
| **Postsynaptic** |  |  |  |
| Absent AChR staining (%) | 16.4±13.4 | 50 ±46.4 | 18.56±16.67 |
| AChR area (µm^2^) | 160.8±82(60) | 65.2±61.3(9) | 70.6±54.4(40) ^***^ |
| AChR perimeter (µm) | 240.4±96(60) | 156.6±124.3(9) | 203.2±134.8(40) |
| Endplate area (µm^2^) | 381.5±158.5(60) | 219.6±96.3(9) | 585.6±462.7(40) ^#^ |
| Endplate perimeter (µm) | 89.6±17.4(60) | 81.5±35(9) | 177.3±249.7(40) |
| Endplate diameter (µm) | 39.8±23.4(60) | 47.2±24.7(9) | 55.9±25.0(40) ^**^ |
| **Derived Variables** |  |  |  |
| **Presynaptic** |  |  |  |
| Average length of branches (µm) | 4.5±2.0(72) | 3.8±1.8(28) | 2.9±1.1(49) ^*^ |
| Complexity | 4.5±0.5(72) | 3.7±0.9(28) ^**^ | 4.4±0.9(49) ^#^ |
| **Postsynaptic** |  |  |  |
| Average area of AChR clusters (µm^2^) | 131.5±97.9(59) | 48.2±70.6(9) | 11.2±10.7(40) ^****^ |
| Fragmentation | 0.2±0.3(61) | 0.6±0.4(9) | 0.8±0.1(40) ^****^ |
| Compactness (%) | 42.5±14(60) | 29.1±18.5(9) | 14.2±10.6(40) ^****^ |
| Overlap (%) | 64.6±15.3(60) | 37.1±30.3(9) | 11.9±14.3(40) ^****^ |
| Area of synaptic contact (µm^2^) | 101.4±51.7(60) | 25.5±37.6(9) ^*^ | 10.8±14.8(40) ^****^ |
| **Associated Nerve Variables** |  |  |  |
| Axon diameter (µm) | 2.1±0.8(47) | 1.7±0.5(26) ^**^ | 2.1±0.7(43) |
| Number of axonal inputs | 1.0±0.1(55) | 1.1±0.3(27) | 1.0±0.1(46) |

Initial analysis of NMJ structure in the soleus revealed no sex related differences in variables and therefore data has been combined for males and females. Analysis was performed using previously published method ^27^. Data presented as mean ± s.d. (number of NMJs analysed). Most variables were not normally distributed and were analysed using Kruskal-Wallis test followed by Dunn’s multiple comparisons test. WT n=7, *Agrn^nmf380^* (Mot) n=3, *Agrn^nmf380^* (3B2) n=5

^*^Significantly different from WT, ^#^significantly different from Mot. ^*/#^P<0.05, ^**/##^P<0.005, ^***/###^P<0.001, ^****/####^P<0.0001.

**Supplementary Table 6: Morphological variables of NMJs from WT and *ColQ^-/-^* mice.**

| **Variable** | **WT** | ***ColQ^-/-^* (Mot)** | ***ColQ^-/-^* (3B2)** |
| --- | --- | --- | --- |
| **Core Variables** |  |  |  |
| **Presynaptic** |  |  |  |
| Absent Synaptophysin staining (%) | 0 | 0 | 0 |
| Nerve terminal area (µm^2^) | 154.3±76.7 (122) | 49.8±35.3(124) ^****^ | 28.7±18.7(127) ^****/#^ |
| Nerve terminal perimeter (µm) | 168.6±80.9(122) | 70.2±42.9(124) ^****^ | 44.3±27.9(127) ^****/#^ |
| Number of terminal branches | 19.3±11.8(122) | 12.3±8.3(124) ^***^ | 1.0±8.2(127) ^****/#^ |
| Number of branch points | 12.3±7.7(122) | 3.5±2.9(124) ^****^ | 1.9±1.9(127) ^****^ |
| Total length branches (µm) | 77.5±42.0(122) | 26.2±17.3(124) ^****^ | 16.3±11.3(126) ^****/#^ |
| **Postsynaptic** |  |  |  |
| Absent AChR staining (%) | 0 | 0 | 0 |
| AChR area (µm^2^) | 160.4±84.8(122) | 64.6±38.5(122) ^***^ | 33.4±29.4(127) ^****/###^ |
| AChR perimeter (µm) | 174.6±91.4(122) | 97.2±53.3(122) ^***^ | 64.8±43.9(127) ^****/##^ |
| Endplate area (µm^2^) | 315.0±167.3(122) | 154.8±99.4(122) ^****^ | 102.8±99.0(127) ^****/#^ |
| Endplate perimeter (µm) | 79.2±22.6(122) | 57.3±22.5(122) ^***^ | 45.2±21.9(127) ^****/#^ |
| Endplate diameter (µm) | 32.7±16.7(122) | 22.7±12.7(122) ^**^ | 17.7±11.7(127) ^****/##^ |
| **Derived Variables** |  |  |  |
| **Presynaptic** |  |  |  |
| Average length of branches (µm) | 4.7±3.3(122) | 2.3±1.2(124) ^****^ | 2.3±1.4(126) ^****^ |
| Complexity | 4.0±0.8(120) | 2.9±0.8(114) ^****^ | 2.4±0.8(97) ^****/####^ |
| **Postsynaptic** |  |  |  |
| Average area of AChR clusters (µm^2^) | 117.4±74.0(122) | 27.0±25.1(122) ^***^ | 10.0±11.5(127) ^****/###^ |
| Fragmentation | 0.0±0.3(122) | 0.0±0.9(122) ^***^ | 0.7±0.3(127) ^****/#^ |
| Compactness (%) | 52.5±12.9(122) | 44.9±12.5(122) ^*^ | 34.3±12.1(127) ^****/####^ |
| Overlap (%) | 63.2±14.8(122) | 38.3±15.0(122) ^****^ | 35.4±18.6(124) ^****^ |
| Area of synaptic contact (µm^2^) | 102.5±57.6(122) | 24.6±17.5(122) ^***^ | 11.5±9.8(124) ^****/##^ |
| **Associated Nerve Variables** |  |  |  |
| Axon diameter (µm) | 2.4±1.1(106) | 1.4±0.7(110) ^****^ | 1.3±0.7(113) ^****^ |
| Number of axonal inputs | 1.0±0.0(107) | 1.0±0.1(96) | 1.0±0(94) |

Initial analysis of NMJ structure in the soleus revealed no sex related differences in variables and therefore data has been combined for males and females. Analysis was performed using previously published method ^27^. Data presented as mean ± s.d. (number of NMJs analysed). Most variables were not normally distributed and were analysed using Kruskal-Wallis test followed by Dunn’s multiple comparisons test. WT n=7, *Agrn^nmf380^* (Mot) n=3, *Agrn^nmf380^* (3B2) n=5

^*^Significantly different from WT, ^#^significantly different from Mot. ^*/#^P<0.05, ^**/##^P<0.005, ^***/###^P<0.001, ^****/####^P<0.0001.
